# Fatty acid metabolism in neutrophils promotes lung damage and bacterial replication during tuberculosis

**DOI:** 10.1101/2023.10.25.563909

**Authors:** Poornima Sankar, Ramon Bossardi Ramos, Jamie Corro, Mohd Saqib, Tanvir Noor Nafiz, Gunapati Bhargavi, Anil K Ojha, Yi Cai, Selvakumar Subbian, Bibhuti B. Mishra

**Author notes:** Address of correspondence: Bibhuti B Mishra, PhD.

## Abstract

*Mycobacterium tuberculosis* (Mtb) infection triggers a significant influx of neutrophils to the lungs, which is linked to tuberculosis (TB) severity. The mechanism by which Mtb infection induces neutrophillic inflammation remains unclear. Using a clinically relevant and hypervirulent Mtb strain from the W-Beijing family, HN878, we found that genes related to both glycolysis and fatty acid metabolism are upregulated in the lung neutrophils of susceptible mice. Similar effects in gene expression were observed in rabbits, and humans with pulmonary TB compared to healthy controls. Inhibiting glycolysis with 2-deoxy D-glucose (2-DG) exacerbated disease pathology, while fatty acid oxidation (FAO) inhibitor Etomoxir (ETO) improved outcomes by reducing weight loss, immunopathology, and bacterial replication within neutrophils in genetically susceptible mice. Notably, ETO reduced neutrophil production in the bone marrow and their recruitment to the lungs. ETO specifically restrained the recruitment of Ly6G^low/dim^ immature neutrophil population, which is elevated during disease progression and harbors the bulk of bacilli. In a transwell setup, we demonstrated that ETO dose-dependently inhibited neutrophil chemotaxis towards infected macrophages. In summary, our research highlights the crucial role of fatty acid metabolism in regulating neutrophilic inflammation during TB and provides a rationale for targeting immunometabolism of neutrophils for potential TB treatment.

## INTRODUCTION

Neutrophils, the most abundant white blood cells in the human body, play crucial roles in protective immunity against various pathogens[1,2]. Recent studies, both experimental and clinical, focusing on active pulmonary tuberculosis (TB), have revealed a substantial influx of neutrophils into lung tissues[3–5]. An increase in neutrophil counts and a high neutrophil-to-lymphocyte ratio serve as distinguishing factors between TB patients and healthy individuals who test positive in tuberculin skin tests. Among TB patients, an extensive neutrophilic response signifies the severity of TB and, notably, is associated with damage to the pulmonary architecture[6–8]. In tissue biopsies obtained from pulmonary TB patients, neutrophils are the dominant cell types[9]. Unique transcript signatures of active TB, primarily consisting of blood neutrophil-specific type I IFN-inducible genes, have been discovered[10]. These signatures can accurately identify patients with active disease as distinct from those with latent tuberculosis infection (LTBI). Moreover, cross-species signature specific to neutrophils, capable of distinguishing hosts likely to progress to active disease from those who will contain the infection have been reported [11]. Additionally, the presence of neutrophils and the neutrophil granule protein S100A9 in TB lesions, both in nonhuman primates and humans, is indicative of individuals with latent infections likely to progress to active disease [12,13]. Notably, elevated neutrophil counts in the blood are predictive of increased mortality risk from TB[14]. Collectively, these reports firmly establish neutrophils as key pathological mediators in TB. While the specific role and clinical significance of neutrophils in TB remain a subject of debate, several studies have highlighted a strong connection between the development of TB and the infiltration of tissues by neutrophils.

Neutrophils are the primary phagocytes initially recruited from the bloodstream to the lungs during Mtb infection [15]. Individuals in contact with pulmonary TB patients show higher peripheral blood neutrophil counts and a lower likelihood of Mtb infection. *In vitro* studies have demonstrated that neutrophils can significantly inhibit Mtb growth, albeit with variability among donors. They achieve this by causing oxidative damage to the bacteria [16]. Neutrophil-depleted blood exhibits reduced capacity to control Mtb infection *ex vivo* [17]. Overall, these findings also emphasize the importance of neutrophils in the innate antimicrobial immune response against Mtb.

Neutrophils in TB have been overlooked recently, in part due to their limited presence in commonly used murine models like C57BL/6 and BALB/c mice, which are relatively resistant to Mtb infection [18]. In a classic low-dose aerosol Mtb infection, neutrophils were only observed in the lungs within the first 14 to 21 days post-infection and were scarcely detected in later stages [4,19]. These lung lesions primarily consisted of macrophages, T- and B-lymphocytes, lacking the necrotic granulomas seen in human patients[20].[21]. Interestingly, neutrophil depletion in these mice did not impact disease outcome. The decline in neutrophil presence in these lesions was attributed to IFN-g-induced Nos2 expression, which curbs uncontrolled IL-1-mediated neutrophil influx following the onset of adaptive immunity [4]. In contrast, mice deficient in Nos2, IFN-g, or T-/B-lymphocytes succumbed to disease, developing TB lesions enriched in neutrophils and high bacterial burden. Notably, in mouse strains highly susceptible to TB, such as C3HeB/FeJ and I/St mice, neutrophils constituted a significant portion of the cellular infiltrates at the infection site [18]. In these strains, neutrophils were associated with necrotic tissue damage and early mortality [22]. Depleting neutrophils in Mtb-infected I/St mice improved wasting disease, reduced mycobacterial burden, mitigated pathology, and enhanced survival [5,23].

Neutrophil depletion studies in animal models of TB have suggested a pro-bacterial role of these cells during Mtb infection[3,5,24]. Interestingly, neutrophils affect granuloma outcomes in non-human primates, as depleting neutrophils from granulomas with high bacterial burden reduces replication in those granulomas, while depleting neutrophils from granulomas with low load increased the number of bacteria per granuloma [25]. These results suggest that the degree to which neutrophils contribute to pathogenesis varies along a spectrum at the individual granuloma level. Thus, studying the functions of neutrophils at different stages of the disease may provide a deeper understanding of TB pathogenesis.

In this study, we utilized a replication reporter of the hypervirulent Mtb strain HN878 to establish a murine infection model[26] where we investigated the critical role of neutrophil metabolism in immunity against TB. By targeting glucose metabolism with 2-deoxy D-glucose (2-DG) and fatty acid metabolism with a panel of inhibitors directed at various steps in mitochondrial fatty acid oxidation (FAO), we demonstrate that inhibiting the FAO pathway enhances the host’s ability to restrict intracellular bacterial replication. Treatment of infected mice with Etomoxir (ETO), prevented weight loss, immunopathology, and reduced bacterial replication in the lungs of TB susceptible mice. The inhibition of FAO significantly impacted the phagocytic uptake of Mtb by neutrophils, subsequently affecting the pathogen’s replication within these phagocytes. Notably, ETO decreased the recruitment of neutrophils to the lungs. In a transwell chemotaxis assay, we demonstrated that ETO dose-dependently inhibited neutrophil migration towards infected macrophages. Overall, our results show that fatty acid metabolism plays a vital role in controlling neutrophilic inflammation and suggest that targeting the way neutrophils use energy could be a potential therapeutic approach against TB.

## Results

### Neutrophils are the major immune cells harboring bacteria during TB disease

One of the defining features of tuberculosis (TB) is the inflammation that damages tissues, leading to a decline in lung function and a significant increase in mortality associated with the disease[27]. However, the cellular basis of the inflammatory environment in TB lung that is associated with disease susceptibility is poorly defined. We have previously reported that unchecked neutrophil influx to the lung is associated with susceptibility to the lineage 4 strain Mtb H37Rv infection[4,5,28]. To determine whether similar inflammatory environments develop during Mtb HN878 infection, a clinically relevant hypervirulent strain belonging to lineage 2 of W-Beijing isolates, we challenged mice with approximately 100 colony-forming units (CFU) of aerosolized Mtb. To investigate the presence of bacteria and bacterial replication dynamics, we employed a dual reporter strain of Mtb, Mtb HN878 smyc’-mCherry: SSB2-GFP. These bacteria consistently express mCherry and possess GFP fused to the single strand binding protein (SSB-2) that congregates at the replication fork forming GFP labeled foci. Consequently, replicating bacteria express both mCherry with GFP foci, allowing them to be distinguished from non-replicating bacteria using fluorescence microscopy [29,30] and flow cytometry. Utilizing this dual reporter strain, we infected C57BL/6 (referred as BL/6) mice that are recognized as relatively resistant to TB compared to C3HeB/FeJ (referred to as C3H), inducible nitric oxide synthase deficient (*iNos^-/-^*) and mice lacking the interleukin-1 receptor (*Il1r1^-/-^*). At day 29 pi, total and infected neutrophils were analyzed in the single cell preparation of the lungs of these animals (**Fig. 1A**). There was an augmented influx of neutrophils into the lungs of the susceptible mouse strains compared to BL/6. Moreover, number of neutrophils harboring the Mtb bacilli (infected neutrophils) significantly elevated in the lungs of these susceptible animals compared to BL/6 (**Supplementary Fig. 1A, B**) suggesting a direct correlation of neutrophils and bacterial growth within these cells with TB susceptibility to Mtb HN878 infection.

**Figure 1:**
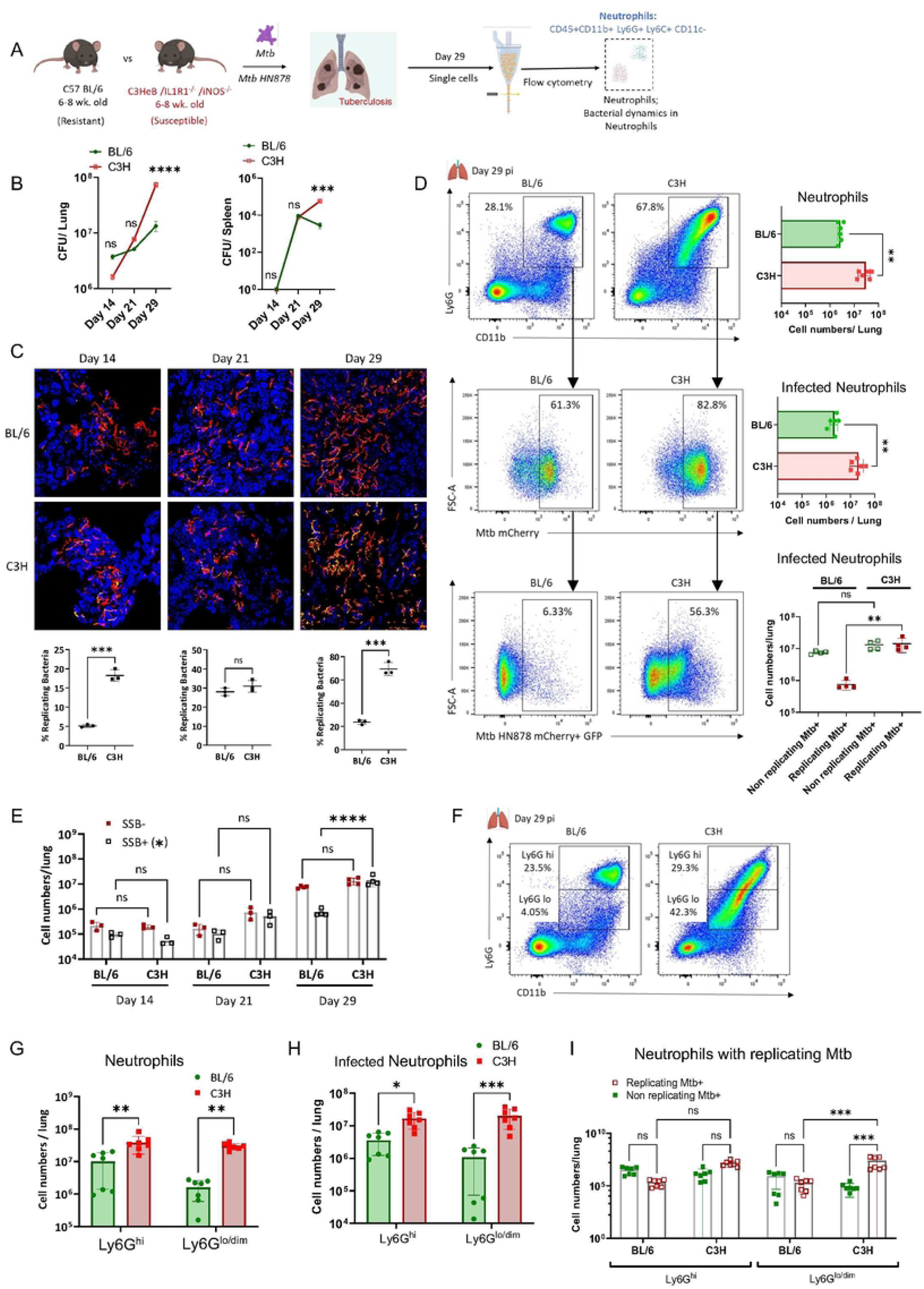
Neutrophils are the major immune subset harboring replicating bacteria during TB disease. (A) Experimental Outline: Six to eight-week-old mice are infected with approximately 100 CFU (colony-forming units) of Mtb HN878 replication reporter (S-myc’-mCherry:SSB2-GFP) via aerosol. Neutrophils (CD45+ CD11b+ Ly6G+ Ly6C+ CD 11c-) were analyzed by flow cytometry on day 29 post-infection (pi). (B) CFU Count: Evaluation of CFU count in lung and spleen homogenates from infected BL/6 and C3H mice at day 14, 21, and 29 pi. (C) Representative Confocal Images: Displaying lung sections from infected BL/6 and C3H mice exhibiting replicating bacteria depicted by GFP foci on mCherry+ bacilli. Additionally, quantification of SSB2-GFP Foci was conducted at indicated time points post-infection. (D) Neutrophil Analysis: - Top Panel: Representative flow cytometry plots and quantification of the number of infiltrated neutrophils in BL/6 and C3H mouse lungs at day 29 pi. - Middle Panel: Representative flow cytometry plots and enumeration of infected neutrophils in BL/6 and C3H mouse lungs at day 29 pi. - Bottom Panel: Representative flow cytometry plots and count of infected neutrophils with replicating bacteria in BL/6 and C3H mouse lungs at day 29 pi. (E) Number of infected lung neutrophils containing replicating bacteria at day 14, 21, and 29 pi. (F) Representative Flow cytometric plots and (G) absolute count of Ly6G^hi^ and Ly6G^l0/dim^ neutrophils in the BL/6 and C3H mouse lungs at day 29 pi. (H) Enumeration of infected Ly6G^hi^ and Ly6G^l0/dim^ neutrophils in BL/6 and C3H mouse lungs at day 29 pi. (I) Count of neutrophils harboring replicating and non-replicating bacteria in Ly6G^hi^ and Ly6G^lo/dim^ neutrophils from BL/6 and C3H lungs at day 29 pi. N=3-6 mice/group- pooled from 2-3 experiments; Error bars = Mean ± SD. **(B, E and I)** 2-way ANOVA; **(C, D)** Unpaired t-test; **(G, H)** Ordinary one-way ANOVA. Statistical significance was calculated by One-way/two-way ANOVA with Tukey’s multiple comparison tests. *: p<0.05; **: p<0.01; ***: p<0.001; ****: p<0.0001.

Next, we analyzed the lung environment of C3H mice at different time points post infection for the neutrophil influx, bacterial replication within neutrophils to ascertain that bacterial replication in neutrophils is a marker of susceptibility. Although differences were not initially apparent at the earlier time points (day 14 and 21 post-infection), susceptible C3H mice exhibited increased neutrophil accumulation and the presence of Mtb-infected neutrophils in their lungs at a later time point, specifically on day 29 post-infection, compared to BL/6 (**Supplementary Fig. 1C, D**). This observation was supported by the increasing bacterial burden in the lung and spleen of C3H mice compared to BL/6 (**Fig. 1B**). We subsequently investigated whether the higher bacterial load in C3H mice was due to enhanced bacterial replication. By enumerating the number of SSB2-GFP foci in mCherry^+ve^ Mtb in the 5μm lung sections, we identified an increased number of replicating Mtb in the C3H lungs compared to BL/6 (**Fig. 1C**).

We have previously shown that neutrophils serve as a permissive niche for Mtb in C3H mice [3,5,19,24]. To examine if the increased bacterial growth in the lungs is a consequence of higher intracellular replication, we probed the replication dynamics of Mtb in neutrophils from infected BL/6 and C3HeB mice at different time points post infection. While neutrophils in BL/6 mice harbor fewer replicating Mtb (SSB-GFP+), approximately half of the lung neutrophils from C3H mice hosted replicating bacteria (**Fig. 1D, lower panel**). No significant difference was observed in the number of infected neutrophils with replicating bacteria at day 14 and 21 pi. However, this difference in Mtb replication was more significant at day 29 pi which corresponded with neutrophil accumulation (**Fig. 1E**).

Recently, bacteria permissive neutrophil population, Ly6G^low/dim^ has been reported in murine models of Mtb H37Rv infection [19,31]. Notably, these immature granulocytes accumulate in the lung as infection progresses and is more abundant in the lung of genetically susceptible mice[19]. To examine if Mtb HN878 infection also recruits these neutrophils to the lung, we compared the neutrophils from BL/6 and C3H mice for their Ly6G expression. We observed an elevated number of Ly6G^lo/dim^ neutrophils in the C3H lungs at day 29 pi (**Fig. 1F, G**). Notably, there was an increase in the infected Ly6G^lo/dim^ neutrophils carrying replicating Mtb in C3H mice compared to BL/6 at day 29 pi (**Fig. 1H and I**). These results support previous observations made in *iNos^-/-^* mice[19] and further indicate that accumulation of Ly6G^low^ immature neutrophils provides a replication conducive cellular niche for Mtb.

In the rabbit model of pulmonary Mtb infection, the outcome of the infection depends on the specific Mtb strain used [32]. When rabbits are infected with Mtb HN878, they develop a progressive disease characterized by severe inflammation and lung pathology marked by necrotic, caseating granulomas. Some of these granulomas progress to form cavities[33,34]. On the other hand, infection with Mtb CDC1551 results in infection control and the spontaneous establishment of latency after initial limited bacterial growth and disease pathology. It’s worth noting that the varying pathological manifestations observed in rabbit lungs infected with these two clinical Mtb isolates closely resemble the spectrum of active TB observed in humans [35,36]. Therefore, we reason that the rabbit model serves as a valuable tool for determining the relationship between neutrophil influx and disease progression.

We evaluated neutrophil influx in the lungs of rabbits infected with Mtb HN878 or CDC1551 at day 14 and 29 pi through comparative pathology. Similar to mice, we observed increased necrotic lesions, heightened neutrophil influx and presence of Acid-fast bacilli (AFB) identified by Ziehl-Nielsen staining of the lungs of rabbits infected with Mtb HN878 (**Supplementary Fig. 1E, G**). In contrast, rabbits infected with CDC1551 had more lymphocytes and histiocytes in their lungs (**Supplementary Fig. 1F**), and nearly undetectable AFB in the lung sections (not shown), which is consistent with a more controlled infection induced by this clinical isolate. These findings collectively suggest that Mtb HN878 infection triggers a robust influx of neutrophils, with these cells being the primary infected cells in the lung and support Mtb replication.

### Expression of metabolism associated genes are increased in neutrophils during TB

Since neutrophils from different host environments were not only quantitatively different, but they also varied in terms of their ability to harbor replicating Mtb, we sought to determine the transcriptome profile of these cells isolated from BL/6 and C3H mice lung. Neutrophils were extracted from the lungs of BL/6 and C3H mice following Mtb infection, on day 27 pi and conducted bulk mRNA sequencing. Principal Component Analysis (PCA) unveiled significant transcriptomic differences among the lung neutrophils (**Fig. 2A**). Through differential gene expression (DEG) analysis, we identified 452 upregulated and 1003 downregulated genes in C3HeB mouse lung neutrophils when compared to those from BL/6 mice (**Fig. 2B**).

**Figure 2:**
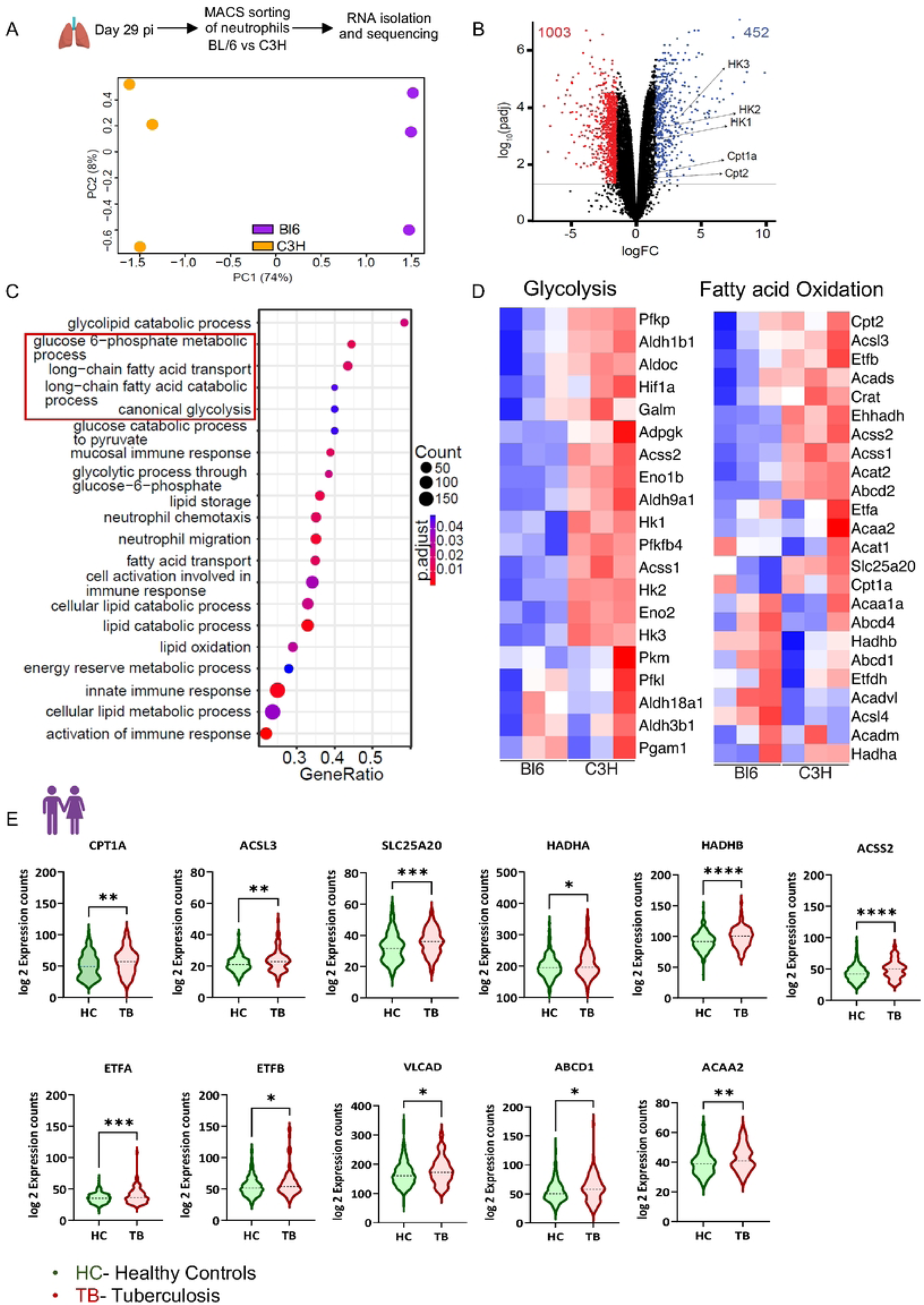
Metabolism associated genes are upregulated in neutrophils from susceptible mice. 6-8-week-old C57BL/6 (BL/6) and C3HeB/FeJ (C3H) mice were infected with approximately 100 CFU of Mycobacterium tuberculosis (Mtb) HN878 replication reporter via aerosol. Neutrophils were isolated from the lung using magnet-associated cell sorting (MACS) at day 27 post-infection (pi) and subjected to mRNA sequencing. (A) Principal component analysis of the RNA sequencing dataset demonstrates the distinct distribution of gene sets in neutrophils from BL/6 and C3H mice. (B) The volcano plot illustrates the differential expression of genes in the resistant BL/6 and C3H neutrophils. (C) Graphs depict selected pathways derived from the top 50 differentially expressed genes between BL/6 and C3H mouse neutrophils. (D) Heatmaps exhibit selected genes from glycolysis and fatty acid oxidation pathways in BL/6 and C3H mouse neutrophils; with a sample size of n=3 mice per group. (E) Log2 fold expression counts of genes involved in the Fatty Acid Oxidation pathway were evaluated through high-throughput RNA sequencing in whole blood cells obtained from TB patients and healthy controls participating in a household contact study. Error bars = Mean ± SD. Statistical significance was calculated by unpaired t-test. *: p<0.05; **: p<0.01; ***: p<0.001; ****: p<0.0001.

Subsequently, we performed gene ontology analysis of these DEGs to delineate the pathways that were up-and downregulated in lung neutrophils. Notably, we observed an upregulation in genes associated with metabolic pathways, including glycolysis, fatty acid transport, uptake, and catabolism in the neutrophils from C3H mice, compared to their resistant BL/6 counterparts (**Fig. 2C**). Specifically, all genes essential for regulating glycolysis and β-oxidation of fatty acids exhibited increased expression in C3H mouse neutrophils from the infected mice (**Fig. 2D, Supplementary Fig. 2A**). We then examined the metabolic gene signature in the previously published genome wide microarray dataset for the Mtb HN878 and CDC1551 infected rabbit lungs. There was an overall increase in the expression of metabolism associated genes in the Mtb HN878 as early as 2 weeks pi compared to CDC1551, linking these gene clusters in inflammatory disease and host susceptibility to TB (**Supplementary Fig. 3B**).

To check the relevance of these gene signatures in human TB, we analyzed the publicly available databases for transcriptomic signatures in the peripheral blood cells (PBMCs) from a group of 98 TB patients and 314 healthy controls participating in a household contact study (GSE94438). This analysis revealed an elevated expression of genes associated with the fatty acid oxidation (FAO) pathway. These genes included CPT1A, ACSL3, SLC25A20, HADHA, HADHB, ACSS2, ETFA, ETFB, VLCAD,ABCD1, and ACAA2 mirroring the patterns observed in mouse lung neutrophils (**Fig. 2E**). Other genes related to fatty acid and glucose metabolism were also altered in PBMCs of TB patients compared to healthy controls (**Supplementary Fig. 3C, D**). Together, these cross-species gene signatures related to glucose and fatty acid metabolism suggest that metabolic programming during Mtb infection could be linked to TB pathogenesis.

To explore whether there were differences in glucose and fatty acid uptake during Mtb infection, we labeled both Mtb-HN878 (smyc’-mCherry) infected and uninfected naïve bone marrow neutrophils with 2-Deoxy-2-[(7-nitro-2,1,3-benzoxadiazol-4-yl)amino]-D-glucose (2-NBDG), a fluorescent glucose analog and BODIPY™ FL C_16_ (4,4-Difluoro-5,7-Dimethyl-4-Bora-3a,4a-Diaza-*s*-Indacene-3-Hexadecanoic Acid (C16-BODIPY), a fluorescent fatty acid analog. Subsequently, we assessed their uptake by flow cytometry as described previously [29]. Our results revealed that fatty acid uptake increased during Mtb infection while glucose uptake was inhibited (**Supplementary Fig. 3A-C**). Next, we sorted neutrophils from the BM of naïve mice, infected *ex vivo* with Mtb HN878-mCherry at an MOI 3 and compared the glucose and fatty acid uptake efficiency of Mtb infected (Mtb+) and uninfected neutrophils (Mtb-) though all neutrophils have been exposed to Mtb. Although infected and uninfected neutrophils have comparable glucose uptake efficiency, Mtb-infected neutrophils significantly take up more fatty acid than Mtb primed yet uninfected neutrophils (**Supplementary Fig. 3D, E**). These findings indicate that fatty acid metabolism may be preferred over glucose for energy during neutrophil’s response to Mtb infection.

### Fatty acid metabolism of neutrophils regulates uptake and replication of Mtb

Genes associated with metabolic pathways are significantly upregulated in susceptible C3H mouse neutrophils when compared to their resistant BL/6 counterparts. To investigate the impact of inhibiting the two primary energy generation pathways, glycolysis, and fatty acid metabolism, on the response of neutrophils to Mtb, we treated *ex-vivo* cultures of sorted naïve bone marrow neutrophils with two inhibitors: 2 deoxy D-glucose (2-DG), a glucose analog blocking glycolysis and Etomoxir (ETO), an inhibitor of fatty acid oxidation (FAO) through its irreversible inhibitory effects on the carnitine palmitoyl-transferase 1a (CPT1a)-mediated β-oxidation. Concurrently, we infected them with Mtb HN878, replication reporter strain (**Fig 3A**).

**Figure 3:**
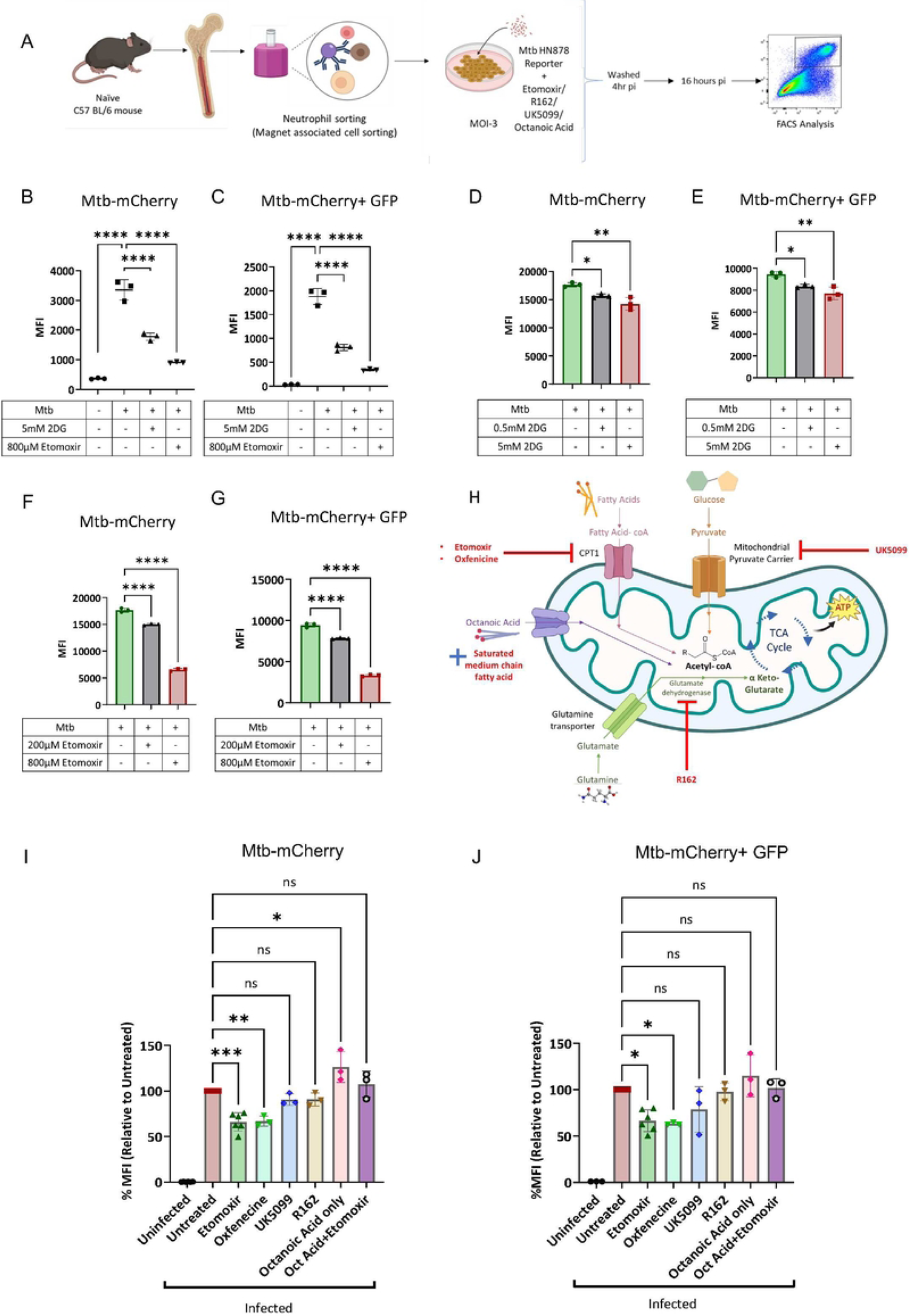
Bacterial uptake and replication by neutrophils are regulated by mitochondrial fatty acid metabolism. (A) Experimental Overview: Neutrophils isolated from the bone marrow of 6-8-week-old naive BL/6 mice by Magnetic sorting. Subsequently, they were infected with Mtb HN878 replication reporter at MOI-3. After a 4-hour post-infection (pi), cells were washed to remove extracellular bacteria. Bacterial uptake (4hr pi) and survival (18hr pi) were analyzed by flow cytometry with and without indicated concentrations of 2-DG and Etomoxir (see methods). (B) The Mean Fluorescence Intensity (MFI) of Mtb HN878 mCherry and (C) MFI of Mtb HN878 SSB2-GFP were evaluated using flow cytometry at 4 hours pi. (D-E) Neutrophils were subjected to treatments with either 0.5mM or 5mM 2DG, followed by an assessment of the MFI of both bacteria (mCherry) and replicating bacteria (GFP) using flow cytometry at 18 hours pi. (F-G) Another set of neutrophils was treated with either 200pM or 800pM Etomoxir. The subsequent MFI of Mtb-HN878-mCherry and Mtb-SSB2-GFP was analyzed using flow cytometry at 18 hours pi. (H) A visual depiction illustrates the mechanism of action of different small molecule inhibitors targeting mitochondrial fatty acid metabolism. Neutrophils were exposed to either Etomoxir, Oxfenicine, UK5099, R162, or Octanoic acid, with some also receiving Octanoic acid combined with Etomoxir. The assessment of the samples after 18 hours pi included (I) MFI of Mtb HN878 mCherry and (J) MFI of Mtb SSB2 GFP. This was compared to untreated controls. N=3 replicates/group; Error bars= mean ± SD. (B-J) Ordinary one-way ANOVA. Statistical significance was calculated by One-way ANOVA with Tukey’s multiple comparison tests. *: p<0.05; **: p<0.01; ***: p<0.001; ****: p<0.0001.

To evaluate the effect of glycolysis and fatty acid metabolism inhibition on phagocytic uptake of bacteria, we measured the mean fluorescence intensity (MFI) of the total (mCherry) and replicating bacilli (SSB2-GFP foci) at 4 hours pi in neutrophils treated with 2-DG or ETO. Remarkably, blocking these metabolic pathways inhibited bacterial uptake by neutrophils (**Fig 3B, C**). However, after 18 hours pi, there was a dose-dependent decline in the MFI of mCherry and mCherry+GFP+ bailli in both 2-DG- and ETO-treated neutrophils compared to vehicle treated cells (**Fig 3D-G**). These results suggest that glucose and fatty acid metabolic pathways are activated during Mtb infection and targeting them may be a viable strategy to control infection.

To address the possibility that the observed effect was due to the inhibition of the desired metabolic pathways, not a result of direct toxicity to bacteria, we exposed Mtb to 5mM 2-DG or 800 µM ETO in 7H9 broth cultures for 18 hours and then plated the bacteria to enumerate the Colony-Forming Units (CFU). This experiment revealed that 2-DG is indeed toxic to bacteria in culture, suggesting that the reduced bacterial uptake and replication in neutrophils could be attributed to the chemical toxicity directly on the bacteria, though contribution of host glucose metabolism in these processes could not be ruled out. However, no toxicity of ETO on Mtb was apparent in this setting (**Supplementary Fig. 4A**).

To further investigate the impact of modulating fatty acid metabolism on neutrophil’s response to Mtb, we targeted the fatty acid metabolic pathways using chemical inhibitors and assessed both uptake and replication of Mtb after 4h and 18hr pi. Mitochondrial pyruvate transport and amino acid metabolism were targeted using UK5099 and R162, respectively. To rule out any off-target effects of ETO other than fatty acid metabolism, we utilized another CPT1a inhibitor, Oxfenicine, to assess if its effects mirrored those of ETO. We also examined if adding fatty acids could reverse the effects of ETO. In this regard, we supplemented ETO-treatment with octanoic acid, a medium-chain fatty acid (**Fig. 3H**). Oxfenicine exhibited similar effects to ETO, and the addition of octanoic acid alongside ETO reversed its effects. However, treatment of infected neutrophils with UK5099 or R162 did not impact bacterial uptake or replication in neutrophils, suggesting a specific effect related to fatty acid oxidation for generating energy (**Fig. 3J-I**; **Supplementary Fig. 4C, D**). Importantly, none of these inhibitors caused any significant changes in neutrophil viability compared to untreated controls (**Supplementary Fig. 4B**).

Previous studies have suggested a role of fatty acid metabolism in the macrophage antibacterial response[37,38]. Blocking fatty acid metabolism enhances the ability of macrophages to eliminate the laboratory strain H37Rv. To investigate whether ETO had similar effects on macrophages during Mtb HN878 infection, we infected bone marrow-derived macrophages (BMDMs) with the replication reporter of Mtb HN878 and treated them with increasing concentrations of ETO. While we observed a significant decrease in the mean fluorescence intensity (MFI) of mCherry and GFP at 4 hours post-infection, we did not detect any significant differences after 3 days post-infection (pi) (**Supplementary Fig. 4 E-H**). Together, these results support the conclusion that fatty acid oxidation regulates the phagocytic uptake in neutrophils and macrophages while promoting the intracellular replication of Mtb within neutrophils.

### Inhibition of fatty acid oxidation ameliorated TB disease by reducing neutrophil infiltration

Our *in-vitro* experiments have clearly demonstrated a metabolism-induced effect during Mtb infection of neutrophils. Consequently, we sought to investigate whether suppressing glycolysis or fatty acid metabolism in the immunocompetent C3H mice, known for their neutrophil-driven susceptibility to TB (**Fig. 1**), could improve disease outcomes. We infected mice with the Mtb HN878 replication reporter strain and treated them with 250mg/kg of 2-DG or 20mg/kg ETO every alternate day starting at day 21 to day 28 pi. Subsequently, we analyzed these mice at day 29 pi (as depicted in the schematics of **Fig. 4A**). ETO treatment, reduced weight loss (**Fig. 4B**), bacterial burden in the lungs by ∼ 10-fold, as indicated by CFU counts (**Fig. 4C**), and improved lung pathology, characterized by reduced necrotic lesions (**Fig. 4D**) compared to control mice that showed wasting, necrotic lung inflammation, and higher bacterial growth, hallmark features of TB disease. While there was a marginal improvement in weight loss and decline in bacterial load, the area of necrotic lung inflammation worsened in 2-DG treated cohorts suggesting that global inhibition of glycolysis may affect the hosts disease tolerance mechanisms (**Fig. 4B-D**). Next, we employed confocal microscopy to image lung sections from untreated controls, 2DG-treated, and ETO-treated mice, with DAPI counterstaining to examine the presence of mCherry^+ve^ bacilli and evaluate their replication status by quantifying the SSB2-GFP foci across the mCherry^+ve^ rods. We quantified the number of bacteria in different fields of view and plotted the percentage of bacteria with SSB2 Foci (**Fig. 4E**). These image analyses provided additional evidence that ETO-treated mice controlled bacterial replication more effectively than vehicle treated control animals or those treated with 2-DG. Notably, the reduction in bacterial burden and replication in the ETO treated lungs was associated with a significant decrease in the number of total (**Fig. 4E**, **Supplementary Fig. 5A**) and infected neutrophils compared to control mice and those treated with 2-DG (**Fig. 4F**). ETO treatment further reduced the neutrophils harboring replicating bacteria compared to vehicle treated animals (**Fig. 4G, Supplementary Fig 5B**). These results further suggest that inhibiting FAO restrain the Mtb-induced neutrophil influx and consequently bacterial replication within these myeloid cells.

**Figure 4:**
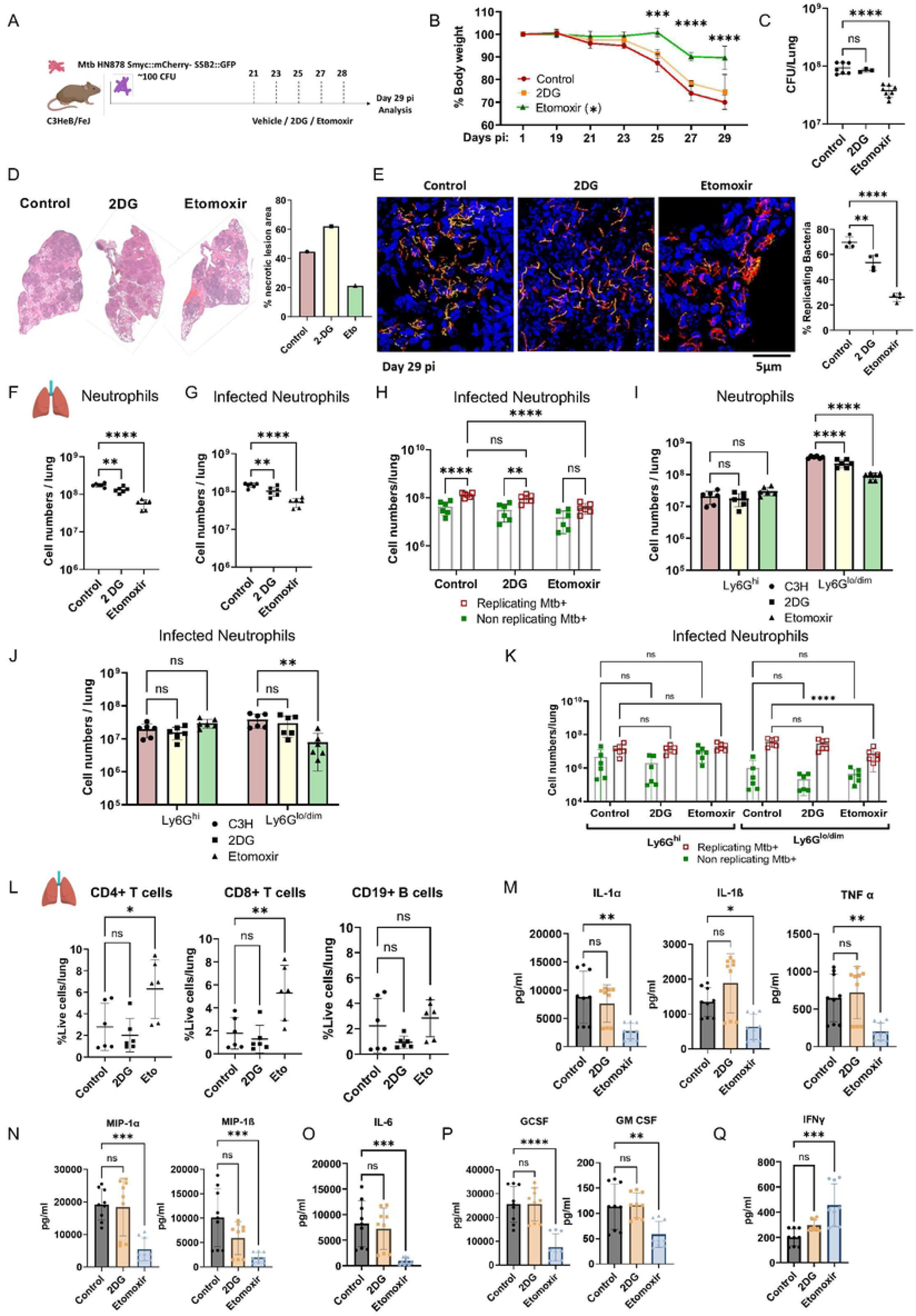
Inhibiting Fatty acid oxidation ameliorates TB Disease *in vivo*. (A) Experimental Overview: 6-8-week-old C3H mice were infected with approximately 100 CFU Mtb HN878 replication reporter via aerosol. They received treatment with either 250 mg/kg 2-DG or 20 mg/kg Etomoxir between day 21 and day 28 and were subsequently analyzed on day 29 post-infection (pi) (B) Graph depicting the percentage of weight loss in untreated, 2-DG-treated, and Etomoxir-treated animals from day 19 to Day 29 pi. Statistics were calculated by comparing the respective time points between Untreated Controls and Etomoxir treated C3H mice. (C) Bacterial burden determined by CFU, in the lungs of untreated control, 2DG-treated, and Etomoxir-treated mice on day 29 pi. (D) Gross histopathology images of untreated control, 2DG-treated, and Etomoxir-treated mouse lungs, alongside a graph illustrating the quantification of % of necrotic lesion area in these lungs. (E) Representative Confocal Images: Displaying lung sections from infected BL/6 and C3H mice exhibiting replicating bacteria depicted by GFP foci on mCherry+ bacilli. Additionally, quantification of SSB2-GFP foci was conducted in different treated samples. (F) Live neutrophils analysis. (G) Analysis of live infected neutrophils. (H) Number of infected neutrophils with replicating and non-replicating bacteria in the lungs of untreated controls, 2DG, and Etomoxir-treated mice at day 29 pi as assessed by flow cytometry. (I) Total count of live Ly6G^hi^ and Ly6G^l0/dim^ neutrophils. (J) Total count of live infected Ly6G^hi^ and Ly6G^l0/dim^ neutrophils. (K) Ly6G^hi^ and Ly6G^to/dim^ neutrophils with replicating and non-replicating bacteria in untreated control, 2DG, and Etomoxir-treated mouse lungs at day 29 pi. (L) Graphs displaying the percentage of live CD4+ T-cells, CD8+ T-cells, and CD19+ B-cells in the lungs of untreated controls, 2DG, and Etomoxir-treated mice at day 29 pi. (M) Cytokine analysis from lung homogenates of untreated controls, 2DG, and Etomoxir-treated mice. Graphs displaying the levels of proinflammatory cytokines, including IL-1 α, IL-1 β, TNF-α. (N) Levels of MIP-1α and MIP-1β. (O) Levels of IL-6. (P) Levels of colony stimulating growth factors G-CSF and GM-CSF and (Q) Protective cytokine IFN-γ. N=6 mice/group- representative of 2 experiments; Error bars= mean ± SD. (B, H-K) Two-way ANOVA; (C, E-G, l-Q) Ordinary One-way ANOVA. * Statistical significance was calculated by One-way ANOVA or two-way ANOVA with Tukey’s multiple comparison tests. *: p<0.05; **: p<0.01; ***: p<0.001, ns-non-significant.

We then analyzed the counts of Ly6G^hi^ and Ly6G^lo/dim^ neutrophil populations in the treated mice, as the latter immature neutrophil populations accumulate in the lung with disease progression and have been linked to TB susceptibility (see **Fig. 1E-G**). Specifically, mice treated with ETO displayed a reduction in the accumulation of Ly6G^lo/dim^ neutrophils (**Fig. 4H**). Additionally, a lower number of infected Ly6G^lo/dim^ neutrophils harboring replicating bacteria were observed in the lungs of ETO-treated mice. In contrast, the infiltration of the mature Ly6G^hi^ cells remained unaffected by ETO treatment, and the bacterial replication within these cells did not show a significant difference between the treated and untreated mice (as shown in **Fig. 4I, J**). Furthermore, ETO treatment led to a decrease in the infiltration of monocytes, macrophages, eosinophils, and plasmacytoid dendritic cells into the lung (**Supplementary Fig. 5C**). Moreover, there was an increased proportion of protective CD4+ and CD8+ T-lymphocytes in ETO-treated mice (**Fig. 4L and Supplementary Fig. 5D**), suggesting that the FAO blockade enhanced protective immunity against TB.

To investigate the effect of ETO treatment on the inflammatory lung environment of C3H mice, we examined 32 different cytokines, chemokines, and growth factors in the lung homogenates. These factors have been associated with the regulation of inflammation, particularly in the context of neutrophil-mediated pathological responses. Consistent with a reduction in lung inflammation, the mice treated with ETO exhibited decreased levels of several pro-inflammatory cytokines, including IL-1α, IL-1β, and TNF. Moreover, chemokines such as MIP-1α, MIP-1β, and MIP-2, known to be involved in monocyte and neutrophil trafficking, were reduced. Additionally, growth factor cytokines G-CSF and GM-CSF, which provide survival signals for neutrophils, were significantly diminished compared to those treated with 2-DG or the vehicle-treated controls (**Fig. 4M-P; Supplementary Fig. 6**). Conversely, we observed elevated levels of the protective cytokine IFN-γ in the lung homogenates of the ETO-treated group, suggesting the creation of a protective host environment through the inhibition of fatty acid metabolism (**Fig. 4Q; Supplementary Fig. 6**). Overall, these findings suggest that fatty acid metabolism promotes heightened neutrophil recruitment to Mtb-infected lungs and increased bacterial replication in TB lesions, particularly within neutrophils, thereby contributing to an increased susceptibility to TB.

### Fatty acid metabolism is crucial for neutrophil production and migration

Previous studies have demonstrated that inhibiting fatty acid metabolism in neutrophils through CPT1a mediates an impact on neutrophil chemotaxis [39]. To investigate whether the inhibition of CPT1a by ETO in neutrophils affects their migration in response to infected macrophages, we conducted a transwell chemotaxis assay. Neutrophils, along with varying concentrations of ETO, were introduced into the upper chamber, and their migration across a collagen coated semi-permeable membrane in response to Mtb-infected macrophages was assessed after an 18-hour incubation period. Post-migration analysis revealed a dose-dependent effect of ETO on neutrophil migration in response to Mtb-infected macrophages (**Fig. 5A**). Thus, it appears that fatty acid metabolism plays a crucial role in the chemotaxis of neutrophils in response to Mtb infection.

**Figure 5:**
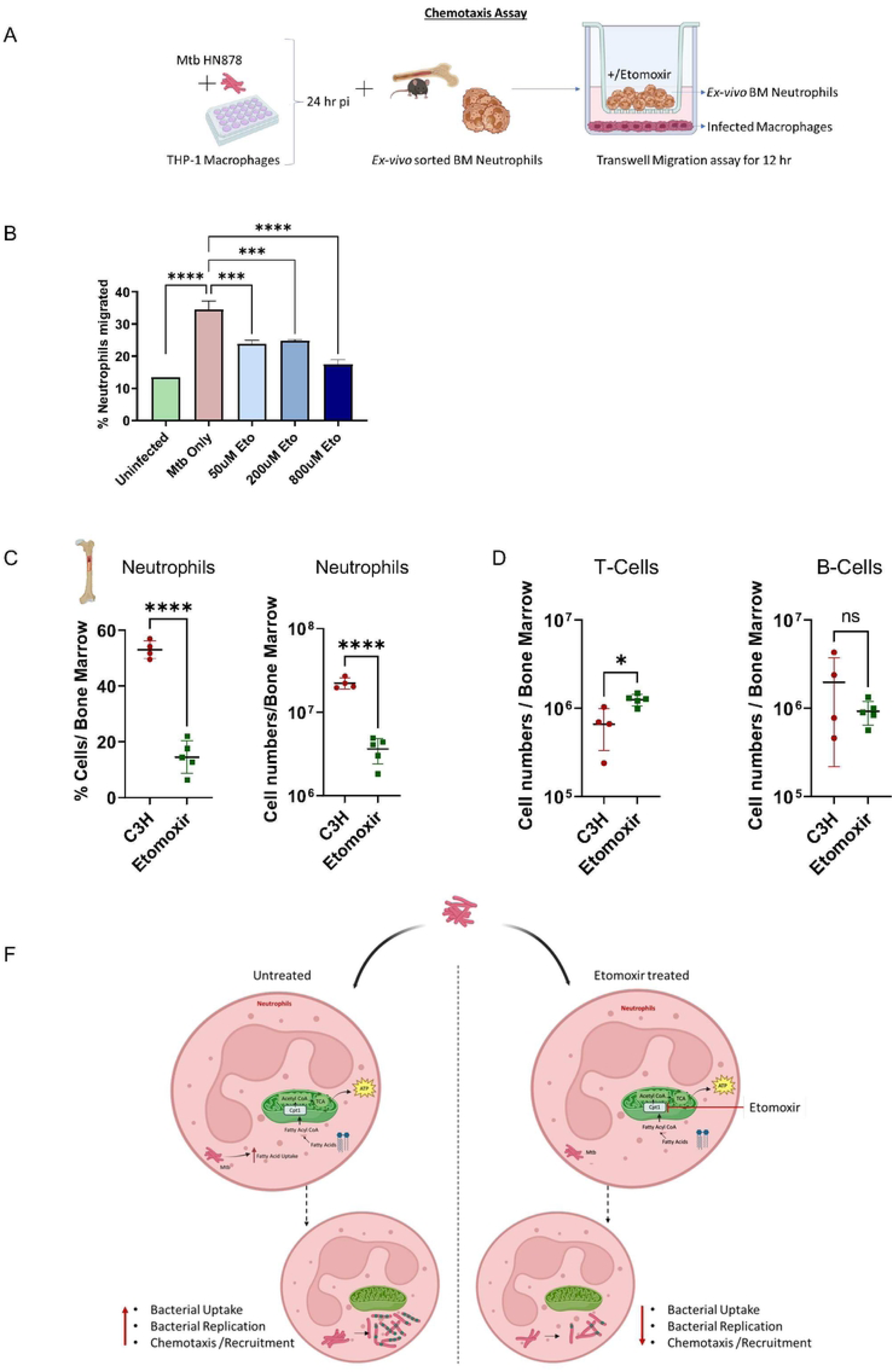
Restriction of fatty acid metabolism *in-vivo* affects bone marrow production of neutrophils. (A) Schematics showing the overview of the transwell migration assay. (B) Macrophages were infected with Mtb strain HN878 for 24 hours. Neutrophils were then treated with varying concentrations of Etomoxir and added in a collagen coated transwell chamber. The migration of neutrophils to the bottom chamber in response to the infected macrophages was quantified. A graph was generated to represent the percentage of neutrophil chemotaxis into the bottom chamber, analyzed by the transwell migration assay. (C) C3H mice were administered 20 mg/kg of Etomoxir, and the bone marrow compartment from the infected mice was analyzed at day 29 post-infection. Graphs showing the percentage and the total number of neutrophils, (D) the total count of T cells (CD11b-CD19-CD3+) and B cells (CD11b-CD3-CD19+) in both untreated control and Etomoxir-treated mice at day 29 post-infection. (E) A model depicting the effect of Fatty acid oxidation blockade on Mtb-neutrophil interactions. N=4-5 mice/group; Error bars= mean ± SD. Statistical significance was calculated by One-way ANOVA with Tukey’s multiple comparison tests (B); Unpaired t-test (C-E). *: p<0.05; **: p<0.01; ***: p<0.001; ***: p<0.001; ns-non-significant.

Considering the impact of fatty acid metabolism on neutrophil migration, we inquired whether the reduced neutrophil influx into the lungs of ETO-treated animals resulted from diminished recruitment or due to decreased synthesis of these cells in the bone marrow. We repeated the ETO treatment experiment as shown in **Fig. 4** in a separate cohort. The control cohorts received phosphate buffer saline (PBS) as vehicle. We assessed the number of neutrophils in the bone marrow compartment of mice treated with ETO by flow cytometry and observed an overall reduction in the percentage and total number of neutrophils compared to vehicle-treated controls (**Fig. 5B, C**). Concurrently, there was an increase in the number of CD3e+ T-lymphocytes in the bone marrow of ETO-treated mice compared to the control cohort (**Fig. 5D**). However, CD19+ B-lymphocytes were not altered by ETO treatment. Collectively, these results demonstrate that fatty acid metabolism may be critical for neutrophil production in the bone marrow and their recruitment to the site of infection (see the model in **Fig. 5E**).

## DISCUSSION

Despite recent advances in the field of immunometabolism, our understanding of neutrophil metabolism remains incomplete. Previous studies have indicated that neutrophils primarily rely on glucose as their primary carbon source for metabolic processes [40]. However, emerging evidence suggests that neutrophils also utilize a variety of other nutrients, including amino acids, carbohydrates, proteins, and lipids, to generate energy [41,42]. Neutrophils frequently encounter immunological environments with limited nutrient availability, necessitating their ability to adapt and employ diverse metabolic pathways to meet the demands of the immune response. For instance, immature neutrophils exhibit a greater dependence on oxidative phosphorylation (OXPHOS) and fatty acid oxidation (FAO) due to their higher mitochondrial content compared to mature neutrophils. In contrast to mature neutrophils, immature neutrophils generate energy by degrading lipid droplets through a process known as lipophagy, subsequently channeling the resulting fatty acids into the tricarboxylic acid (TCA) cycle and OXPHOS [41]. Our findings emphasize the crucial role of fatty acid metabolism in the recruitment of Ly6Glow/dim immature neutrophils to the lungs. However, it is worth noting that inhibiting glucose uptake by 2-DG also affected the recruitment of these cells, suggesting that both fatty acid and glucose metabolic pathways may be necessary for energetics, although the former pathways are preferred. Conversely, FAO blockade did not influence the infiltration of mature Ly6G^hi^ neutrophils, consistent with previous reports in cancer models[43]. Furthermore, the glycolysis inhibitor had no impact on these cells, suggesting that mature neutrophils may utilize alternative energy sources for their effector functions, at least in our *in vivo* model.

Recent studies have shed light on the connection between Mtb pathogenesis and host metabolism [44]. Among the primary cellular niches for Mtb, macrophages play a crucial role. Within the lungs, alveolar macrophages (AMs) and interstitial macrophages (IMs) represent the major populations of infected macrophages, each exhibiting distinct metabolic profiles. AMs, which primarily rely on fatty acid oxidation (FAO), create a permissive environment for Mtb replication. In contrast, glycolytically active IMs limit infection[29]. Mtb enhances its reliance on mitochondrial oxidative metabolism, particularly exogenous fatty acids, and induces the formation of lipid-droplet-filled “foamy” macrophages [28]. These foamy macrophages are often found in the inner layers of TB granulomas. Bacilli can be found in close proximity to intracellular lipid droplets, which are believed to serve as a source of nutrients in the form of cholesterol esters and fatty acids, creating a hospitable niche for the bacterium. Near these foamy macrophages, neutrophils reside in the granulomatous lesions and are exposed to the lipid-rich caseum [5,45]. Neutrophils could also serve as a source of these lipids in TB lesions. However, the metabolic status of neutrophils in TB lesions has not been previously investigated. In this study, we employed chemical inhibitors of mitochondrial β-oxidation of fatty acids, such as ETO, which inhibits FAO by targeting CPT1. CPT1 is essential for the entry of long-chain fatty acids into the mitochondrial matrix. ETO not only reduced bacterial uptake but also significantly decreased the number of replicating bacteria intracellularly. The coupling of fatty acid oxidation and phagocytosis, as well as the mechanisms by which neutrophils control bacterial replication, remain open questions. As observed in macrophages, one potential antimicrobial factor could be mitochondrial ROS (mROS) generated due to impaired electron flow in the electron transport system. mROS may recruit NADPH oxidase complexes to neutrophil phagosomes to restrict bacterial replication as has been reported to be a pivotal antimicrobial mechanism of macrophages [37,46]. Moreover, ROS production has been linked to limiting immunopathology due to TB in murine models by regulating proinflammatory IL-1β production and neutrophil influx mediated by this cytokine [47]. These potential mechanisms warrant further investigation in future studies.

One crucial discovery from our study was the influence of inhibiting FAO on neutrophil chemotaxis. Neutrophil accumulation in the lungs has been associated with tissue damage and exacerbated disease. Therefore, modulating the recruitment of neutrophils to the lungs using ETO could serve as a potential therapeutic strategy for improving host antimicrobial functions. Trimetazidine is an FDA approved drug used for treating cardiovascular complications e.g., angina exhibits nanomolar inhibitory effects on the mitochondrial metabolism. TMZ targets the long-chain 3-ketoacyl-CoA thiolase activity of the hydroxyacyl-coenzyme A (CoA) dehydrogenase trifunctional multienzyme complex subunit beta (HADHB) that catalyzes the final step of β-oxidation of fatty acids. Such host-directed therapies (HDTs) would ideally complement the use of direct antimicrobial agents, potentially shortening treatment regimens. While the *in vivo* impact of FAO inhibition by ETO was impressive in reducing Mtb replication in tissues, especially within neutrophils, it’s worth noting that at the drug concentrations administered in mice, it also affected several other cell types. The ramifications of impairing FAO in various cell types, particularly those belonging to the myeloid and lymphoid lineages, represent an area for future investigation as it will unravel new mechanisms of pathogenesis. Previous studies have indicated off-target effects of ETO at concentrations exceeding 10 µM, which were attributed to the disruption of acetyl CoA homeostasis. Concentrations surpassing 200µM have been demonstrated to impair IL-4 production from macrophages, thereby affecting M2-polarization of these cells [48]. However, in our study, we did not observe any significant alteration in IL-4 and IL-13 cytokines in the lungs of mice treated with ETO (**Supplementary Fig. 6**). Nonetheless, we could not dismiss the possibility of other potential off-target effects. Therefore, future investigations employing genetic models are necessary to establish the role of FAO in neutrophil’s bioenergetics in regulating TB pathogenesis.

Unlike FAO inhibition, the inhibition of glycolysis by 2-DG had a moderate effect on bacterial replication but exhibited a significant exacerbation in tissue pathology **(Fig.4D)**. 2-DG, an inhibitor of glucose uptake, completely inhibits glycolysis and affects the linked pentose phosphate pathway (PPP) and the TCA cycle, which utilizes pyruvate, an end-product of glycolysis, as a substrate. Therefore, the observed increase in necrotic foci in the lung could be a result of the toxicity caused by the global inhibition of glycolysis in cell types that play a protective role during TB immunity. Indeed, a recent report has highlighted the protective role of glycolysis in myeloid cells for TB control[49]. These authors demonstrated that myeloid cell-specific deletion of lactate dehydrogenase subunit A enhances susceptibility to TB. Hence, the seemingly toxic effect of 2-DG treatment *in vivo* could be a consequence of the loss of these protective effects. Therefore, an experimental approach that selectively targets glycolysis in neutrophils would provide new insights into the role of glucose metabolism in their functions *in vivo* during TB.

In summary, our work highlights the role of mitochondrial fatty acid metabolism in the response of neutrophils to Mtb infection. We discovered that neutrophils prefer fatty acids over glucose during infection, and the fatty acid oxidation pathway is particularly used by immature neutrophils for recruitment to the site of infection. This suggests that targeting the way neutrophils use energy could be a potential approach for TB treatment.

## Materials and Methods

### Mice

C57BL/6 (Strain #:000664), C3HeB/FeJ (Strain #:000658), IL1R1^-/-^ (Strain #:003245), iNos^-/-^ (Strain #:002596) mice were purchased from The Jackson Laboratory. 6–8-week-old mice were used in this study. They were bred and maintained under Specific Pathogen Free conditions at Albany Medical College. All mouse studies were conducted in accordance with protocols approved by the AMC Institutional Animal Care and Use Committee (IACUC) (Animal Care User Protocol Number ACUP-21-04003). Care was taken to minimize pain and suffering in Mtb-infected mice.

### Bacterial strains

Hypervirulent *Mycobacterium tuberculosis* (Mtb) HN878 were used throughout this study. Mtb HN878 strains transformed with fluorescence reporters smyc’ mCherry: SSB2 GFP or only smyc’: mCherry maintained under Hygromycin B resistance. Bacteria were cultured in Middlebrook 7H9 media with OADC supplement, 0.05% Tween 20 and 0.5% v/v Glycerol and 50 µg/ml Hygromycin B in a shaking incubator at 37 °C for 5-7 days up to log phase for bacterial infections. Strains were maintained in 20% Glycerol in −80°C.

### Mouse infections

Single cell suspension of Mtb HN878 strains was prepared in Phosphate Buffer Saline containing 0.05% Tween 80 (PBST). Clumps were dissociated by passing through 18 Gauge and 25 Gauge needles and ∼100CFU bacteria were used for infection through aerosol route using a Glas-Col inhalation exposure system, Terre Haute, IN. Infection was assessed by Colony Forming Unit (CFU) enumeration in serially diluted lung and spleen homogenates from infected mice at day 29 dpi by plating on 7H10 Agar plates containing 0.5% v/v Glycerol and OADC enrichment. Colony counting was done in plates incubated at 37°C after 3 weeks.

### *In-vivo* treatment of animals with 2DG and Etomoxir

C3HeB/FeJ mice infected with Mtb HN878 were treated with 2-deoxy glucose (2DG) (250mg/kg) (Cat#111980250-Thermo Fisher) or Etomoxir (20mg/kg) (Cat# 11969-Cayman Chemical) via intraperitoneal route or oral gavage. Percentage weight loss was calculated throughout the course. Mice were euthanized at Day 29 pi and assessed for CFU count, histopathology, cytokines, and immune cells.

### Neutrophil isolation and *in-vitro* infection

Bones from naïve C57 BL/6 mice were flushed with complete DMEM media (DMEM with Sodium Pyruvate, Sodium Bicarbonate, HEPES and 10% FBS). Extracted bone marrows were washed, passed through 18 Gauge needles and then 70µM cell strainers to make single cell suspensions. Red blood cells were lysed by ACK lysis. From these cells, neutrophils were isolated by negative sorting by magnetic selection using the Mojosort neutrophil isolation kit according to the suggested protocol (Cat# 480058-BioLegend). Purity was checked by flow cytometry using CD11b and Ly6G surface staining. Purified neutrophils were counted and cultured in complete DMEM at 37 °C with 5% CO_2._ For *in-vitro* infection, Mtb single cell suspension was prepared as above in complete DMEM and added onto neutrophils at the specified Multiplicity of Infection (MOI). 4 hours pi, cells were washed with fresh culture media to remove extracellular bacteria and they were analyzed by flow cytometry at 18 hours pi.

### Bone Marrow derived Macrophage generation and *in-vitro* infection

Naïve C57BL/6 mouse bones were flushed to isolate bone marrow cells. Single cell suspension of bone marrow cells was made as described above. Following ACK lysis, these cells were cultured for 3-5 days DMEM media containing L929 conditioning media, Sodium Pyruvate, Sodium Bicarbonate, HEPES and 10% FBS. When the bone marrow cells have differentiated into macrophages, the media was changed to complete DMEM and infected at an MOI of 3 with Mtb HN878 dual reporter. Cells were either stained after 4hr or washed to remove extracellular bacteria and stained 3 days pi.

### Flow Cytometry

Lung tissues were harvested from infected mice at stated time points and collected in cold PBS. These tissues were chopped and digested at 37 °C in a Collagenase Type IV (150U/ml) (Cat 17104019-Gibco) and DNase I (60U/ml) (Cat# 10104159001 Roche-Sigma Aldrich) cocktail. Post digestion, the suspension was passed through 70µm cell strainers and ACK lysed to get single cell suspensions that were used for further staining. All subsequent steps were done in cells resuspended in FACS Buffer (PBS+0.5% BSA). Fc-Block CD16/32 (Cat# 156604-BioLegend) was used to restrict non-specific antibody binding. Surface staining was done with directly conjugated antibodies at 4°C in the dark for 30 minutes. The cells were then fixed with Fixation Buffer (Cat# 420801-Bio legend) according to manufacturer’s instructions and analyzed on BD Symphony flow cytometer. All analyses were done in FlowJo v10. Dead cells were excluded with eFluor 780 conjugated fixable viability dye (Cat# 65-0864-14 eBioscience) staining and further gating to analyze various populations was done.

### Immunofluorescence microscopy

Lung tissues from infected mice were fixed in 10% Buffer Formalin and were transferred sequentially to PBS, 15% Sucrose in PBS and 30% Sucrose in PBS and OCT embedded. These blocks were cut at 5 µm thickness and mounted on glass slides for further staining. For replicating bacteria visualization in lung sections, sections were washed for 5 minutes, twice with PBS-0.5% v/v Tween 20 (wash buffer) and once with PBS. The sections were then mounted with Prolong Gold Antifade reagent containing DAPI. All slides were imaged on the Series 200 Dragonfly confocal microscope (Oxford Instruments). The images were deconvoluted using Auto Quant (Media Cybernetics) and analyzed on ImageJ. For quantification, SSB2 foci, as seen by GFP staining were enumerated and the percentage of mCherry^+ve^ bacteria with SSB2-GFP foci is represented.

### Histopathology

Formalin fixed infected lung tissues were paraffin embedded. 5 µm thick sections were cut and counterstained with hematoxylin and eosin (H&E) to analyze pathophysiology. H&E staining was performed at the histopathology core at Albany Medical College and were obtained on the NanoZoomer 2.0 RS Hamamatsu slide scanner and analyzed on Image J.

### *In-vitro* treatment of neutrophils

To test *in-vitro* uptake of fatty acids or glucose, 4 hours prior to harvesting infected neutrophils, 25µM C16 BODIPY (Cat# D3821-Invitrogen) or 15µM 2NBDG (Cat# N13195-Invitrogen) was added directly onto the cells. Cells were then stained and processed for flow cytometry. Neutrophils were either untreated or treated throughout the course of infection with 0.5mM 2Deoxy-Glucose (2DG) (Cat#111980250-Thermo Fisher), 5mM 2DG, 200 µM Etomoxir (Cat# 11969-Cayman Chemical), 800 µM Etomoxir, 5mM Oxfenicine (Cat# 33698-Cayman Chemical), 50 µM UK5099 (Cat# 16980-Cayman chemical), 50 µM R162 (Cat# 30922-Cayman Chemical), 100 µM Octanoic Acid (Cat# C2875-Sigma Aldrich) alone or 800 µM Etomoxir with 100 µM Octanoic Acid. Cells were processed as above for flow cytometry.

### Transwell chemotaxis assay

24 well transwell plates with 5 µm pore size were used for this assay. The semipermeable membrane layer was coated with 30 µg/ml collagen in 60% ethanol for at least 4 hours prior to the assay. These trans wells were calibrated with HBSS+ (with Ca^+2^ and Mg^+2^) by washing them twice with HBSS+ and leaving the coated plates in the 24 well plate with 1ml HBSS+ at 37 °C for 1-2 hours. The lower chamber had THP-1 cells (5 × 10^5^ cells/well) in complete DMEM, uninfected or infected with Mtb HN878 at MOI-1, 24 hours prior to the assay. 10^6^ sorted neutrophils either pretreated for an hour with different concentrations of Etomoxir (50 µM, 200 µM, 800 µM) or untreated in 20 µl HBSS+, with or without Etomoxir were added to the top chamber and allowed to migrate for 18 hours. Post migration, the top chambers were discarded. Migrated neutrophils along with trypsinized THP-1 cells were collected, stained, and processed for flow cytometry.

### Cytokine analysis

Mouse lung homogenates were prepared in PBS at Day 29 pi, and total protein concentrations were measured by Pierce BCA protein assay kit according to manufacturer’s instructions. Total protein concentrations were normalized in all samples and sent to Eve Technologies for quantification using the Luminex xMAP technology for multiplexed quantification of 32 Mouse cytokines, chemokines, and growth factors. The multiplexing analysis was performed using the Luminex™ 200 system (Luminex, Austin, TX, USA) by Eve Technologies Corp. (Calgary, Alberta). Thirty-two markers were simultaneously measured in the samples using Eve Technologies’ Mouse Cytokine 32-Plex Discovery Assay® (Millipore Sigma, Burlington, Massachusetts, USA) according to the manufacturer’s protocol. The 32-plex consisted of Eotaxin, G-CSF, GM-CSF, IFNγ, IL-1α, IL-1β, IL-2, IL-3, IL-4, IL-5, IL-6, IL-7, IL-9, IL-10, IL-12(p40), IL-12(p70), IL-13, IL-15, IL-17, IP-10, KC, LIF, LIX, MCP-1, M-CSF, MIG, MIP-1α, MIP-1β, MIP-2, RANTES, TNFα, and VEGF. Assay sensitivities of these markers range from 0.3 – 30.6 pg/mL for the 32-plex. Individual analyte sensitivity values are available in the Millipore Sigma MILLIPLEX® MAP protocol.

### Human patients whole blood transcriptome study

To identify blood biomarkers that might predict TB disease, blood samples from patients diagnosed with TB were collected every 6 months, up to 18 months. People who went on to develop TB disease were considered as TB cases and others who don’t were considered controls (Healthy). The high throughput sequencing data was analyzed on Illumina HiSeq 2000 platform and deposited under GEO GPL11154. Out of 412 subjects, 98 went on to develop TB and 314 remained healthy. The sequencing dataset has been deposited in GEO under series GSE94438.

### Rabbit model of Mtb Infection, Histopathology, AFB staining and Transcriptome analysis

#### 1. Mtb infection

New Zealand White rabbits (specific pathogen-free) of about 2.5 kg (Envigo, USA) were infected with aerosolized Mtb HN878 or Mtb CDC1551 strains using the “nose-only” exposure system (CH Technologies Inc., NJ, USA) to implant ∼3.2-3.5 log_10_ CFU in the lungs (T=0) as previously described[33,50]. At defined time points, rabbits (n=4-6 per time point) were euthanized, and lungs were used for histopathologic examination and RNA isolation. All animal procedures were conducted according to the protocols approved by the Rutgers University IACUC.

#### 2. Rabbit lung histology

Portions of rabbit lungs were randomly cut and fixed in 10% neutral formalin, followed by paraffin-embedding. The formalin-fixed and paraffin-embedded tissue blocks were cut into 5-micron sections and used for staining with hematoxylin & eosin (H&E) or acid-fast staining by Ziehl-Nielsen method (for Mtb). Stained lung sections were analyzed microscopically using Nikon Microphot-FX system with NIS-elements F3.0 software (Nikon Instruments, NY).

#### 3. RNA isolation from rabbit lungs

After the autopsy, random portions of rabbit lungs with or without Mtb infection were immediately treated with TRI Reagent (Molecular Research Center, Cincinnati, OH) and total RNA was isolated, as described previously[34]. RNA was purified using RNeasy kit (Qiagen, CA, USA) and the quality and quantity were measured with Agilent 2100 Bioanalyzer (Agilent Technologies, CA, USA).

#### 4. Genome wide transcriptome analysis of Mtb-infected rabbit lungs

Total RNA from rabbit lungs with or without Mtb infection was subjected to genome wide transcriptome profiling using 4X44k rabbit whole genome microarrays (Agilent Technologies, Santa Clara, CA), as described[51]. Briefly, arrays were hybridized with a mixture of Cy3 or Cy5 labeled cDNA, generated using uninfected or Mtb-infected rabbit lung RNA at each time point. The arrays were washed, scanned and data extracted. Background-corrected, normalized data were analyzed by One way ANOVA using Partek Genomics Suite Ver 6.8 (Partek Inc., St. Louis, MO); an unadjusted p value < 0.05 was used to select significantly differentially expressed genes (SDEG). The list of SDEG was uploaded into Ingenuity Pathway Analysis portal (IPA; Qiagen, CA) as described previously[34], to identify gene networks and pathways impacted by the SDEGs. The microarray data is deposited to Gene Expression Omnibus (accession numbers GSE33094 and GSE39219).

### RNA Library Preparation and Sequencing

Single cell suspensions from Mtb infected mouse lungs were prepared as described above. Mouse neutrophils were sorted by positive selection using anti-Ly6G-APC for 30 minutes at 4°C in the dark followed by secondary staining with anti-APC nanobeads following the APC nanobead kit protocol (Cat# 480090-Bio Legend). Purity was assessed using an anti-mouse GR1 antibody. These Ly6G+ cells were used for RNA isolation. Sorted Ly6G+ cells were stored at −80°C in RLT Plus Buffer (Cat# 1053393-Qiagen). Total RNA isolation was done using RNeasy Plus Mini Kit (Cat# 74134-Qiagen) following the manufacturer’s instructions. RNA samples were quantified using Qubit 2.0 Fluorometer (Thermo Fisher Scientific, Waltham, MA, USA) and RNA integrity was checked with 4200 TapeStation (Agilent Technologies, Palo Alto, CA, USA). rRNA depletion sequencing library was prepared by using QIAGEN FastSelect rRNA HMR Kit (Qiagen, Hilden, Germany). RNA sequencing library preparation uses NEBNext Ultra II RNA Library Preparation Kit for Illumina by following the manufacturer’s recommendations (NEB, Ipswich, MA, USA). Briefly, enriched RNAs are fragmented for 15 minutes at 94 °C. First strand and second strand cDNA are subsequently synthesized. cDNA fragments are end repaired and adenylated at 3’ends, and universal adapters are ligated to cDNA fragments, followed by index addition and library enrichment with limited cycle PCR. Sequencing libraries were validated using the Agilent Tapestation 4200 (Agilent Technologies, Palo Alto, CA, USA), and quantified using Qubit 2.0 Fluorometer (ThermoFisher Scientific, Waltham, MA, USA) as well as by quantitative PCR (KAPA Biosystems, Wilmington, MA, USA). The sequencing libraries were multiplexed and clustered on one flowcell lane. After clustering, the flowcell was loaded on the Illumina HiSeq instrument according to manufacturer’s instructions. The samples were sequenced using a 2×150 Pair-End (PE) configuration. Raw sequence data (.bcl files) generated from Illumina HiSeq was converted into fastq files and de-multiplexed using Illumina bcl2fastq program version 2.20. One mismatch was allowed for index sequence identification.

### RNA Sequencing Data Analysis

After demultiplexing, sequence data was checked for overall quality and yield. Then, raw sequence reads were trimmed to remove possible adapter sequences and nucleotides with poor quality using Trimmomatic v.0.36. The reads were then mapped to the Mus musculus reference genome available on ENSEMBL using Rsubread v1.5.3. Gene counts were quantified by Entrez Gene IDs using featureCounts and Rsubread’s built-in annotation. Gene symbols were provided by NCBI gene annotation. Genes with count-per-million above 0.5 in at least 3 samples were kept in the analysis. Differential expression analysis was performed using limma-voom.

Reads were mapped to Mus musculus genome available on ENSEMBL using the STAR aligner v.2.5.2b). BAM files were generated as a result of this step. Unique gene hit counts were calculated by using feature Counts from the Subread package v.1.5.2. Only unique reads that fell within exon regions were counted. After extraction of gene hit counts, the gene hit counts table was used for downstream differential expression analysis. Using DESeq2, a comparison of gene expression between the groups of samples was performed. The Wald test was used to generate P values and Log2 fold changes. Genes with adjusted P values < 0.05 and absolute log2 fold changes >1 were called as differentially expressed genes for each comparison. Gene ontology analysis was performed on the statistically significant set of genes by implementing the software GeneSCF. The goa_Mus musculus GO list was used to cluster the set of genes based on their biological process and determine their statistical significance. A PCA analysis was performed using the “plotPCA” function within the DESeq2 R package. The plot shows the samples in a 2D plane spanned by their first two principal components. The top 500 genes, selected by highest row variance, were used to generate the plot. The bulk HUVEC RNA-seq data obtained in this publication have been deposited in NCBI’s Gene Expression Omnibus and are accessible through GEO Series accession number GSE244230.

### Statistics

Statistical differences among the indicated groups were analyzed by unpaired two tailed Student’s *t-*test or one way Analysis of Variance (ANOVA) using Tukey’s multiple comparison tests. All statistical analyses were done using Graph Pad Prism 9 software. A p value of <0.05 was considered significant. The *n* numbers and other significant values are indicated in the figures and figure legends (*: p<0.05; **: p<0.01; ***: p<0.001).

## Supplementary Materials

**Supplementary Figure 1:**
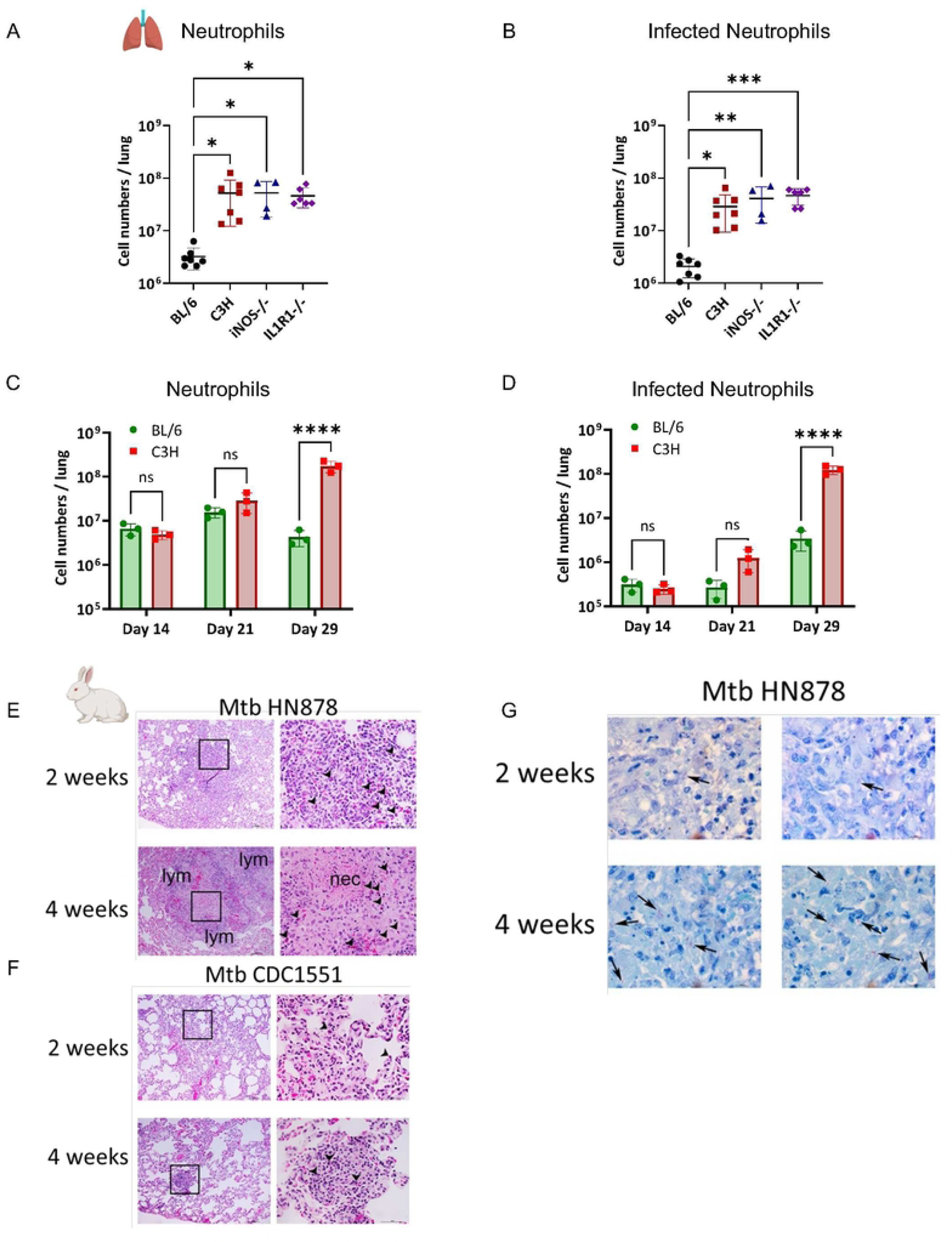
Neutrophils accumulate in mouse lungs with increasing susceptibility to TB. 6-8-week-old mice were infected with Mtb HN878 mCherry SSB2-GFP and evaluated at day 29 post-infection (pi), unless indicated otherwise. (A) Total numbers of live neutrophils in BL/6, C3H, iNos^-/-^ and IL1RT^-/-^ mouse lungs at Day 29 pi. (B) Total number of infected neutrophils in the lungs of BL/6, C3H, iNos^-/-^, and ILIRT^-/-^ mice at day 29 pi. (C) Number of live neutrophils in the lungs of infected BL/6 and C3H mice at day 14, 21, and 29 pi. (D) Number of live infected neutrophils at day 14, 21, and 29 pi. (E) (H) Immunohistochemistry evaluation of rabbit lungs infected with Mtb HN878 at 2 weeks and 4 weeks pi. (I) Assessment of rabbit lungs infected with Mtb CDC1551 at 2 weeks and 4 weeks pi. Square boxes on the left panels of (H) and (I) indicate the magnified area shown on the right. Arrows in the right panels indicate neutrophils, necrosis, and lymphocytes. (F) Acid-Fast Staining of Mtb HN878 from infected rabbit lungs at 2 weeks and 4 weeks pi. N=3-6 mice/group- pooled from 2-3 experiments; Error bars= mean ± SD. **(A, B)** Ordinary one-way ANOVA; (C-E) 2-way ANOVA; Statistical significance was calculated by One-way/two-way ANOVA with Tukey’s multiple comparison tests. *: p<0.05; **: p<0.01; ***: p<0.001; ****: p<0.0001.

**Supplementary Figure 2:**
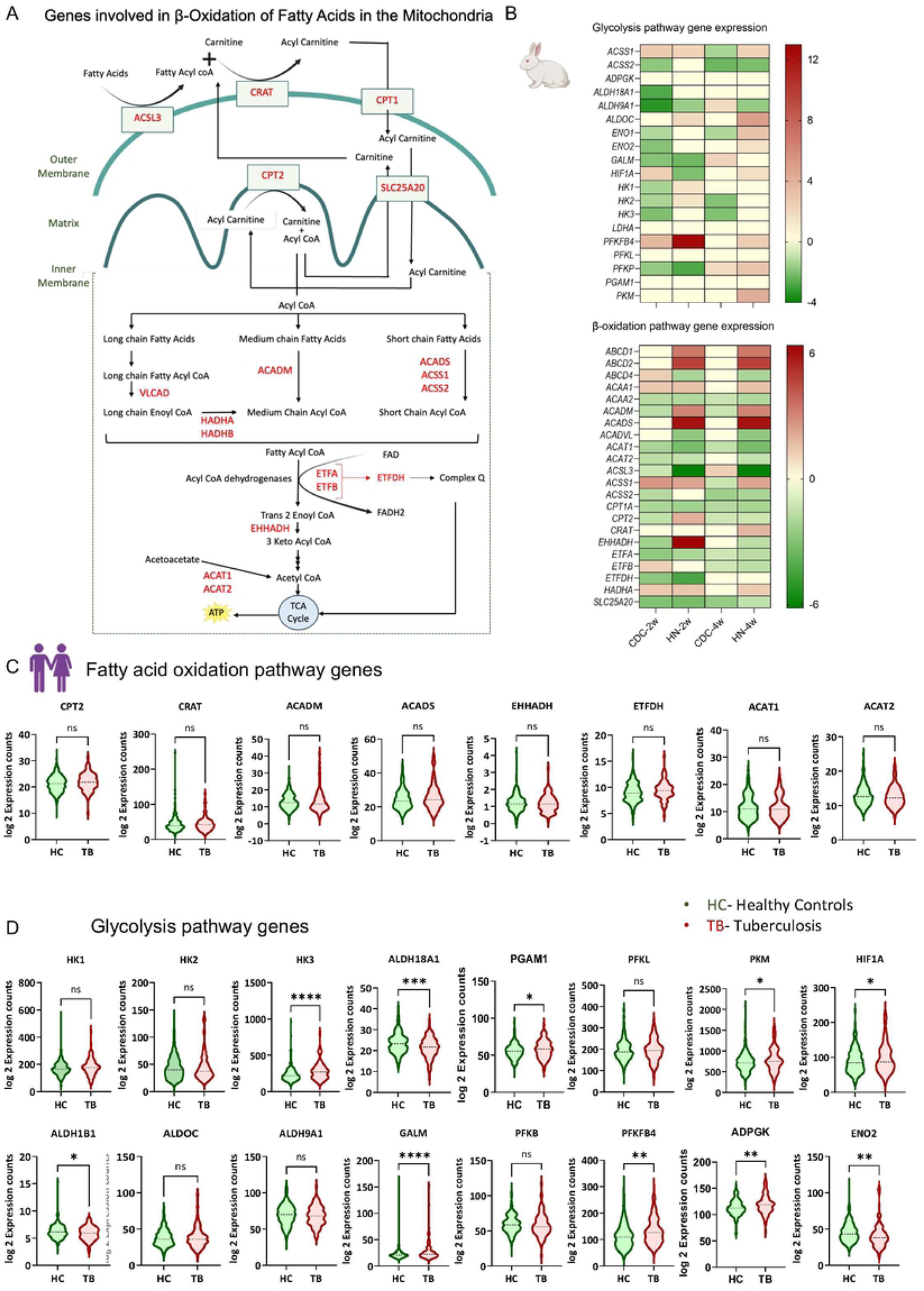
Transcriptome analysis shows an increased expression signature of Glycolysis and Fatty acid oxidation pathway genes. (A) Features schematics highlighting the various genes linked with fatty acid metabolism (highlighted in red) as expressed in murine neutrophils from Mtb HN878 infected mice. (B) Heatmaps were utilized to illustrate the differentially expressed genes related to glycolysis and p-oxidation pathways in the rabbit lungs at 2 weeks and 4 weeks post-infection (pi) with Mtb CDC1551 and Mtb HN878. The determination of gene expression was accomplished through genome-wide microarray analysis. (C) Graphs displaying Iog2 expression counts for specific genes involved in Fatty acid oxidation, while (D) highlights the expression of genes in the glycolysis pathway within the publicly accessible transcriptome dataset GSE94438. This dataset was obtained through RNA sequencing of whole blood cells from a household contact study, involving individuals who developed TB, alongside a control group of healthy individuals. Error bars= mean ± SD. Statistical significance was calculated by unpaired t-test. *: p<0.05; **: p<0.01; ***: p<0.001.

**Supplementary Figure 3:**
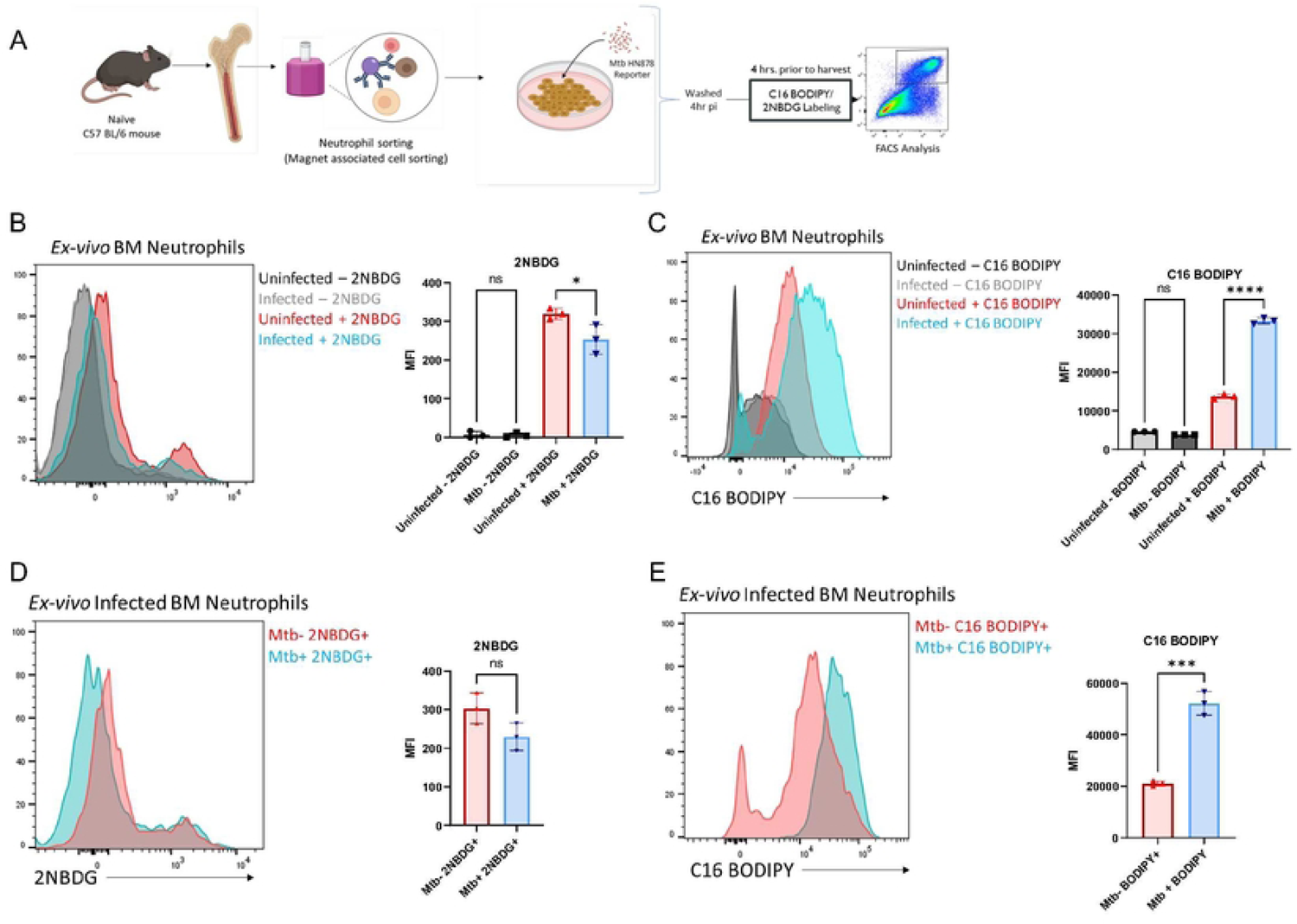
Glucose and fatty acid utilization by neutrophils during Mtb infection. (A) Experiment overview: 6-8-week-old naive BL/6 mouse bone marrow neutrophils were isolated and infected *ex-vivo* with Mtb HN878 s-myc’ mCherry:SSB2-GFP at a multiplicity of infection (MOI) of 3. The cells were washed 4 hours post-infection (pi) to remove extracellular bacteria and then incubated for 18 hours. Four hours before harvest, both infected and uninfected neutrophils were labeled with either C16-BODIPY or 2-NBDG and subsequently analyzed by flow cytometry. (B) Representative flow cytometry histogram plot and graph illustrate the Mean Fluorescence Intensity (MFI) of 2-NBDG in uninfected versus infected neutrophils. (C) Representative histogram and graph depict the MFI of C16-BODIPY labeling in uninfected versus infected neutrophils. (D) Representative flow cytometry histogram and graph display the MFI of 2-NBDG in neutrophils with and without bacteria when neutrophils were challenged with Mtb HN878 ex vivo. Representative flow cytometry histogram plot and graph demonstrate C16-BODIPY MFI in infected (Mtb+) and uninfected (Mtb-) mouse neutrophils when cells were challenged with Mtb HN878 ex vivo. N=3 replicates/group. Error bars= mean ± SD. **(B, C)** Ordinary one-way ANOVA; **(D, E)** Unpaired t-test. Statistical significance was calculated by One-way ANOVA with Tukey’s multiple comparison tests *: p<0.05; **: p<0.01; ***: p<0.001; ****: p<0.0001.

**Supplementary Figure 4:**
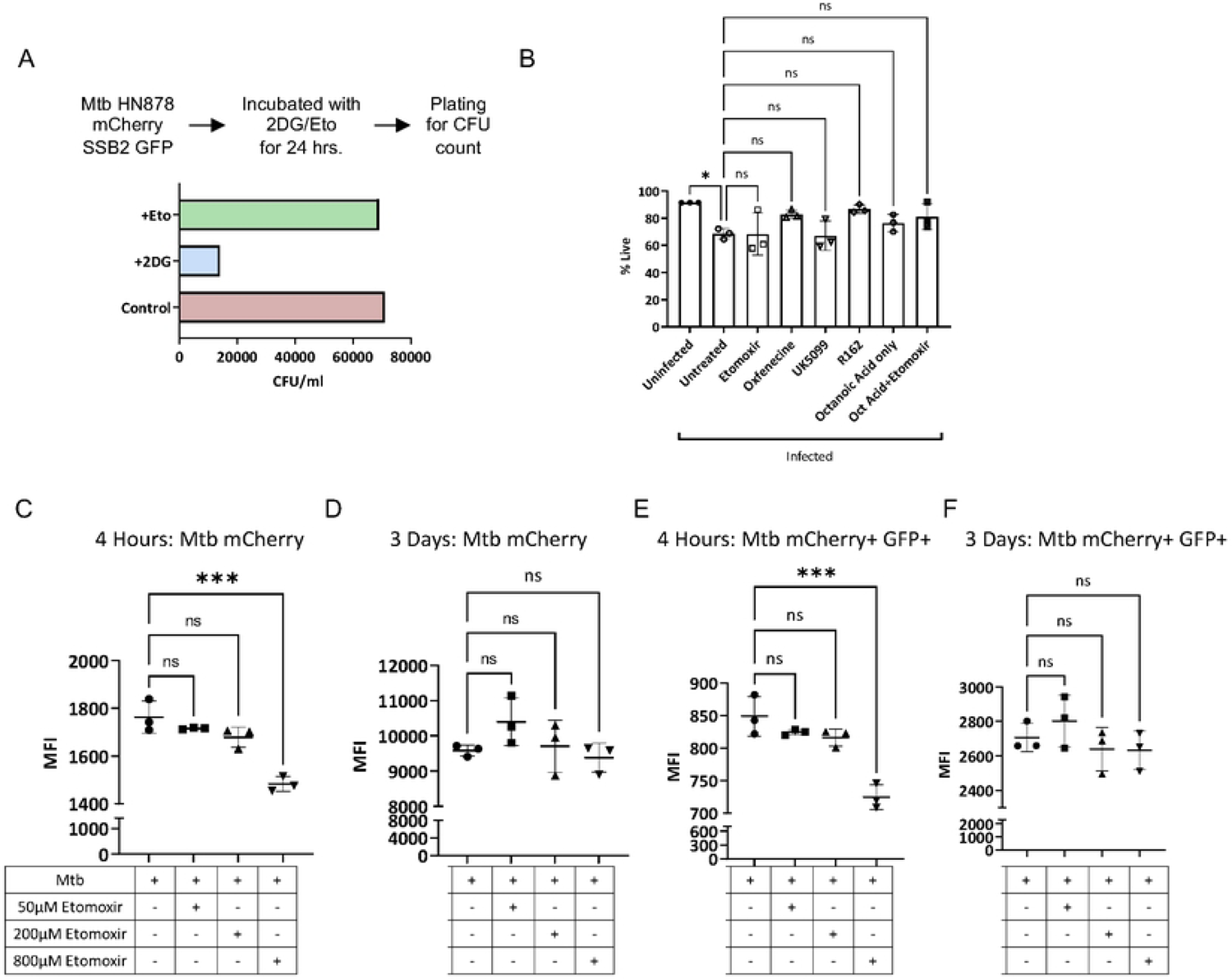
Cytotoxicity of Fatty acid metabolism inhibitors and efefet on macrophage antimycobacterial functions. **(A)** Graph showing CFU of Mtb broth treated with either 5mM 2DG or 800pM Etomoxir. (B) Cell death in neutrophils treated with mitochondrial metabolism inhibitors, including Etomoxir, Oxfenicine, UK5099, R162 or with medium chain fatty acid, Octanoic acid supplement. Bone Marrow Derived Macrophages (BMDM) were made from naive BL/6 mice and were infected at an MOI-3 with Mtb HN878 replication reporter. Graph showing MFI of Mtb HN878 mCherry at **(C)** 4 hours pi and **(D)** 3 days pi. Graph showing MFI of Mtb HN878 SSB2 GFP at **(E)** 4 hours pi and (F) 3 days pi. Compared to untreated controls; n=3 replicates/group, representative of 2 experiments; Error bars= mean ± SD. Statistical significance was calculated by One-way ANOVA with Tukey’s multiple comparison tests *: p<0.05; **: p<0.01; ***: p<0.001, ns-non-significant.

**Supplementary Figure 5:**
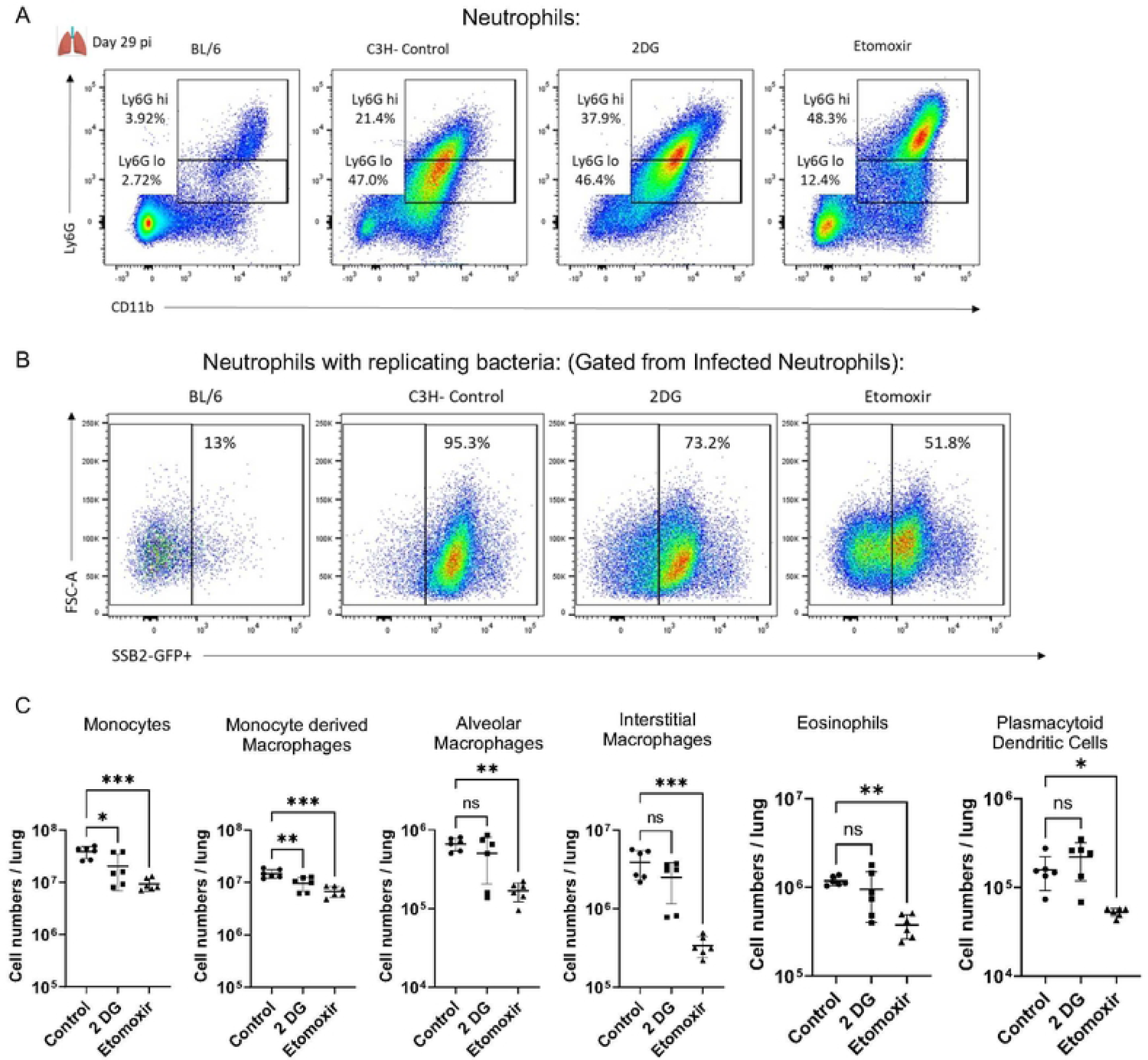
Suppression of Fatty acid metabolism *in-vivo* reduced inflammatory cell influx into diseased mouse lungs. (A) Representative flow cytometric plots from BL/6 and C3H untreated controls, 2-DG-treated, and Etomoxir-treated mice at day 29 post-infection (pi) with Mtb HN878 mCherry SSB2 GFP. (B) Representative flow cytometry plots displaying replicating bacteria in infected neutrophils from BL/6 and C3H untreated controls, 2DG-treated, and Etomoxir-treated mice at Day 29 pi. (C) Graphs illustrating the quantities of live cell populations, including Monocytes (CD45+CD11b+ Ly6G-Ly6C+ SiglecF-F480-), Monocyte-derived Macrophages (CD45+CD11b+ Ly6G-Ly6C+ SiglecF-F480+), Alveolar Macrophages (CD45+CD11b+ Ly6G-Ly6C+ SiglecF -F480+), Interstitial Macrophages (CD45+CD11b+ Ly6G-Ly6C+ SiglecF-), Eosinophils (CD45+CD11b+ Ly6G-Ly6C-SiglecF+), and Plasmacytoid Dendritic Cells (CD11b-Ly6G-Ly6C+ Siglec H+), in untreated controls, 2DG-treated, and Etomoxir-treated mice at day 29 pi. Compared to untreated controls. N=6mice/group- representative of 2 experiments; Error bars= mean ± SD. Statistical significance was calculated by One-way ANOVA with Tukey’s multiple comparison tests. *: p<0.05; **: p<0.01; ***: p<0.001, ns-non-significant.

**Supplementary Figure 6:**
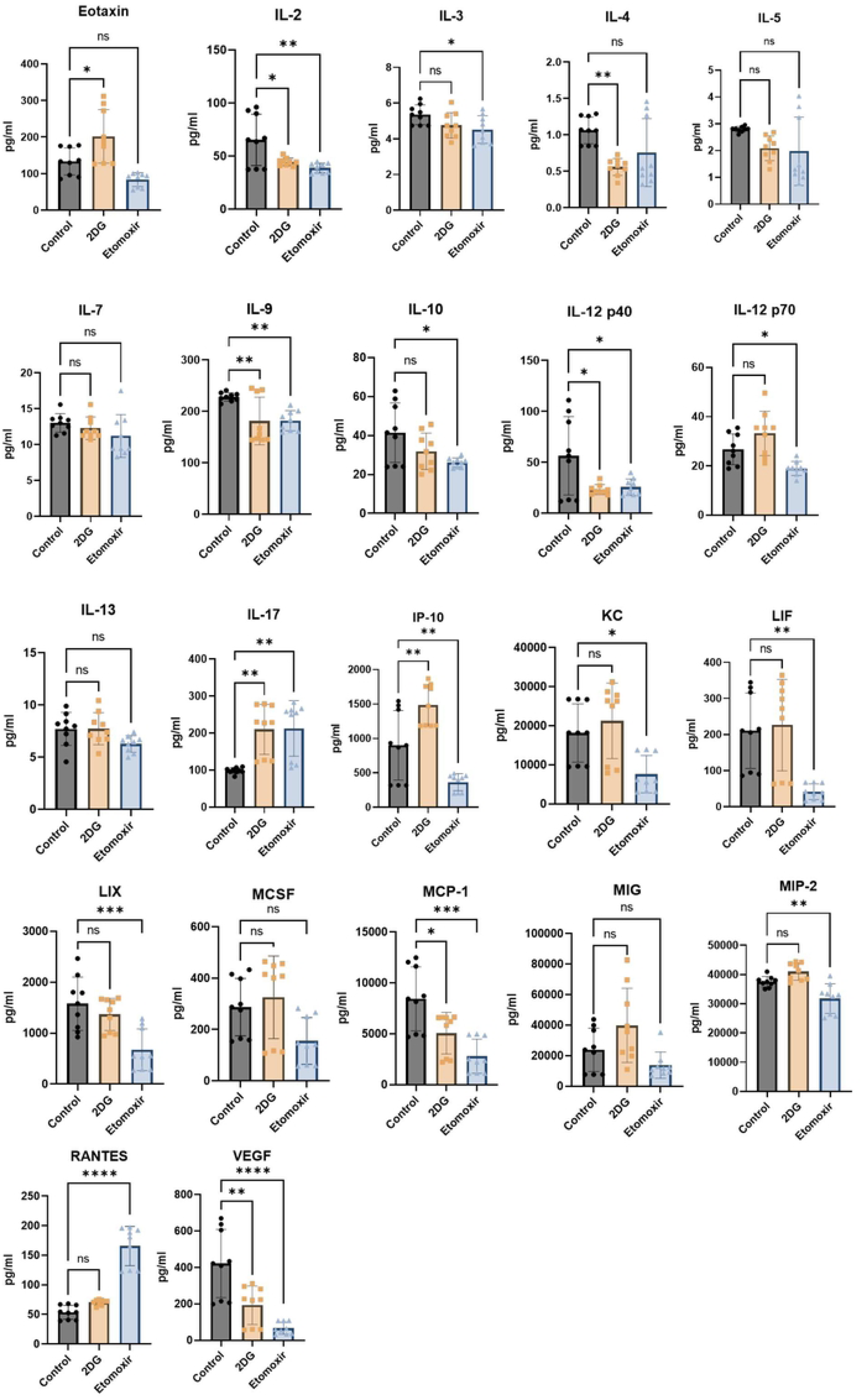
Suppression of Fatty acid metabolism *in-vivo* reduced inflammatory cytokine production in diseased mouse lungs. Cytokine analysis was conducted in lung homogenates from mouse lungs treated with 2DG/Etomoxir. The resulting graphs illustrate the cytokine levels of Eotaxin, IL-2, IL-3, IL-4, IL-5, IL-7, IL-9, IL-10, IL-12p40, IL-12p70, IL-13, IL-17, IP-10, KC, LIF, LIX, MCSF, MCP-1, MIG, MIP-2, RANTES, and VEGF in pg/ml, as compared to the untreated control group. n=6mice/group- representative of two experiments; Error bars= mean ± SD. Statistical significance was calculated by One-way ANOVA with Tukey’s multiple comparison tests. *: p<0.05; **: p<0.01; ***: p<0.001, ns-non-significant.

**Supplementary Table 1:** RAW data used to generate the figures of this manuscript.

## Acknowledgement

We are grateful to the assistance of Christina Peterson and Rebecca Pirri of Pathology and Laboratory Medicine core of Albany Medical College for their help with histopathology and scanning lung tissues. We thank Dr Douglas Cohn, Cindy Vanvorst, Angel Medina-Ramos of Animal Research Facility and Dr. Amit Singh of the BSL3 Management of Albany Medical College for their unwavering support with animal husbandry and care of laboratory animals used in this study.

## Author contributions

Conceptualization and design: BBM and PS. Experiments: PS, BBM, MS, TN. Data analysis and interpretation: BBM, PS, TN. Bioinformatics analysis: RB. Reporter strain generation: AKO. Imaging and data analysis: PS, JC. Human data analysis: YC. Manuscript writing: PS and BBM with input from all co-authors. BBM oversaw the entire study and acquired funding.

## Data and Materials availability

The RNA-seq datasets of mouse neutrophils have been deposited in the NCBI Gene Expression Omnibus database with the accession number GSE244230. Publicly available dataset used in this study to analyze expression of human genes includes GSE94438. RAW dataset used to generate the figures of this manuscript is also included as a supplementary file.

